# Bio-fabricated alginate tumor-like hydrogels to enhance understanding of prostate-specific micro-environments *in vitro*

**DOI:** 10.1101/2025.08.21.671451

**Authors:** K. Al-Husaini, E. Spessot, E. Baena, M. Domingos, A. Tirella

**Affiliations:** Division of Pharmacy and Optometry, School of Biology, Medicine and Health, The University of Manchester, Manchester, UK; Biology Department, College of Science, Sultan Qaboos University, Muscat, Oman; Biotech Center for Biomedical Technologies, Department of Industrial Engineering, University of Trento, Trento, Italy; Cancer Research UK Manchester Institute, The University of Manchester, Manchester, UK; Department of Mechanical and Aerospace Engineering, School of Engineering, Faculty of Science and Engineering & Henry Royce Institute, The University of Manchester, Manchester, UK

**Keywords:** Prostate cancer, 3D *in vitro* model, Alginate, Mechanical properties, 3D printing

## Abstract

Engineered three-dimensional (3D) *in vitro* models are useful tools for closely mimic human tissue-specific tumor microenvironments (TME) and to provide key information on cell-material interactions. In this work we aimed at engineering prostate-specific *in vitro* models to help in discerning specific cell-material interactions at the interface during prostate cancer (PCa) progression, focusing on modelling both biomechanical and biochemical traits of the prostate extracellular matrix (ECM) within PCa progression.

Here, we functionalized alginates and obtained PCa-specific hydrogels for 3D culture and ease evaluation of markers used to assess PCa progression in human prostate cancer cells (i.e., PC-3 cells). Alginate-based hydrogels were modified with laminin-like peptides (i.e., IKVAV, AG73) and tailored in physical and mechanical properties to closely mimic the PCa ECM, with mechanical properties in the range of 2.5-13 kPa (the stiffer value matching advanced/metastatic PCa). To engineer the heterogeneity of advanced PCa, cancer-associated fibroblasts (hTERT PF179T CAF) were selected as stromal cellular component and co/cultured with PC-3 cells. We formulated *prostate bioinks* for extrusion-based bioprinting (EBB) and 3D printed engineered PCa *in vitro* models to study the effect of the microenvironment on the expression of key markers in PC-3 cells, considering the pivotal role of epithelial-to-mesenchymal transition (EMT) in PCa progression.

Cells cultured in prostate-specific hydrogels showed higher cell proliferation and viability, whereas CD44 and Vimentin expression evidenced a higher metastatic potential in PC-3 cells cultured in stiffer and laminin-enriched hydrogels. The selected PCa TMEs used in this work showed PC-3 cells expressing increased levels of Vimentin when co-cultured with CAFs, which also correlates with CD44 expression. Results suggests positive correlations with clinical findings, underlying that tumor biomechanics holds potential for better understand cancer pathobiology and that new 3D *in vitro* models are urged to unveil how ECM traits regulate PCa progression.

Graphical Abstract

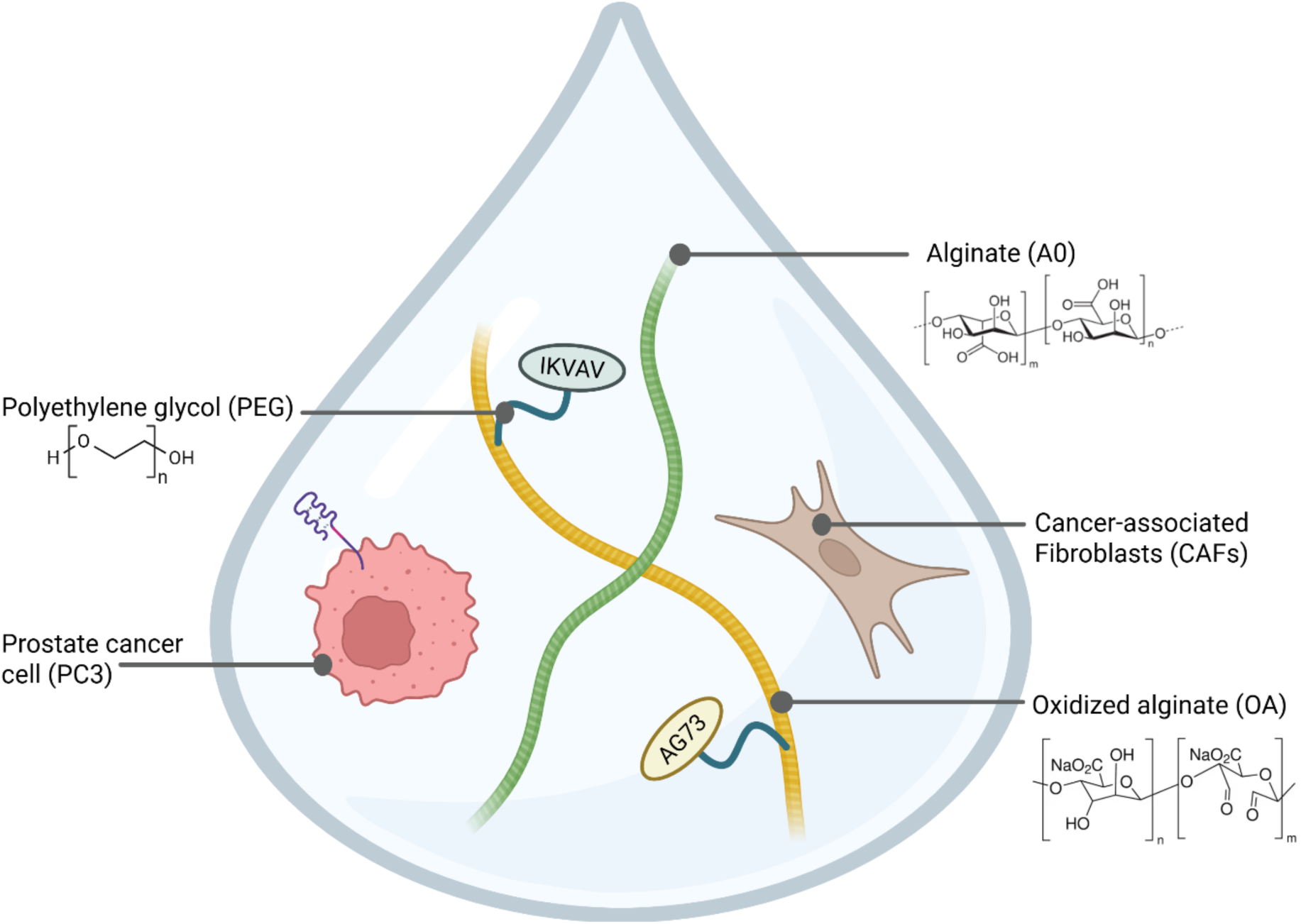

## 1. Introduction

Prostate cancer (PCa) is one of the most commonly diagnosed malignancies and a leading cause of cancer-related deaths worldwide in men ^1^. As in many other solid cancers, the homeostatic disruption caused by cancer cells leads to variations in the composition of the tissue-specific extracellular matrix (ECM) and the interactions with all other cell types (e.g., fibroblasts, macrophages). In PCa, the ECM is a complex network of proteins and polysaccharides that directs tumor progression, metastasis, and drug resistance ^2^; it also provides mechanical support to cells, regulating their behavior both in interactions with other cells and the ECM within the tumor microenvironment (TME) ^3^. Biomechanical and biochemical properties of TME have great impact in PCa progression ^4^, with increased deposition of ECM components, such as collagen, leading to increased mechanical properties (i.e., stiffening) of PCa TME than those of the normal prostate tissue.

What it is known is that the ECM stiffening during PCa progression modulates cells’ phenotype and metastatic potential, yet this remains poorly understood ^5,6^. PCa progression alters ECM components, mainly collagens and laminins, causing prostate tissue stiffening. Among the many prostate-specific ECM components, variation of laminins are being linked to different outcomes inpatients with prostate adenocarcinoma ^7^. Laminins are high molecular weight heterotrimeric glycoproteins composed of three subunits including α-, β-, and γ-subunits, main constituent of the basement membrane with various adhesive and stimulatory functions ^8^. Alterations to the ECM biomechanical properties during PCa progression regulates cell-ECM interactions thus influencing transformation, proliferation, stemness and migration ^8,9^.

Clinically, laminin-α expression is found in different forms (e.g., α1, α3, α5) in prostate ECM, and reported to vary during PCa progression with reported associations between laminins expression and both tumor progression and metastasis ^8,10–12;^ of note, laminin 111 (i.e., α1, β1, and γ1 chains) is the ligand for α6 integrins and detected in elevated levels in grade II and III prostate tumors ^13^. Despite poor knowledge on the correlation between laminins expression in the TME and prognosis scores in clinical settings (e.g., Gleason’s score), differences in laminins expressions in different PCa (e.g., androgen-dependent, castration-resistant prostate cancer) could be linked to changes in laminins and/or integrin expression patterns in cancer cells being predictive for certain clinical profiles.

As such, whilst there is no strong evidence on the correlation between type of PCa and laminin expression in TME, an association between ECM stiffening and laminin expression in patients with prostate adenocarcinoma is reported, starting to link Gleason score and clinical characteristics in PCa progression and metastasis ^7,8^.

Monitoring epithelial–mesenchymal transition (EMT) and cancer stem-like markers (e.g., CD44+) is predictive of the PCa metastatic potential: three-dimensional (3D) culture systems better reflect the *in vivo* situation, enabling new approach methodologies to study cancer cell biology and behavior.

Although, *in vivo* and conventional two-dimensional (2D) *in vitro* models provided fundamental foundation for cancer research, both models faces limitations that challenge their use in advancing cancer research ^14,15^. Typical examples of 3D *in vitro* models are spheroids and organoids developed with different materials ^16^. Hydrogels are often used to support cells (in)growth for cancer research studies, owing the fact that their unique biophysical properties enable to mimic the ECM more effectively ^17^. Among natural polymers used as ECM-mimicking hydrogels, alginate is well-known for its biocompatibility, tunable mechanical properties, and modifications being widely used in 3D *in vitro* models ^18–21^. Extrusion-based bioprinting (EBB) has gained attention for the design and fabrication of 3D *in vitro* models using alginate-based materials, with the advantage of providing precise control over the spatial organization of cells (both cancer and stromal) and the ECM components ^22,23^.

To better understand how the TME impacts on PCa cells progression, invasion and metastasis, this study focused on the manufacturing of engineered prostate-specific 3D *in vitro* models using cancer and stromal human cell lines. Prostate-specific alginate hydrogels were engineered and EBB used to fabricate 3D TMEs, enabling precise control over the biomechanical and biochemical elements of the native prostate TME, as well as controlling the presence of stromal cells (i.e., cancer-associated fibroblasts). The mechanical properties of prostate-specific alginate hydrogels were controlled by varying polymer or crosslinker (CaCl_2_) concentration ^24,25^. Based on our previous studies ^26–28^, oxidized alginate (OA) was prepared and used to include laminin-specific functional groups and by covalently link the aldehyde groups to amino groups as previously shown ^29^. Poly-ethylene glycol (PEG) was used as spacer and linker between alginate and the selected peptides to increase availability of the selected laminin-mimicking adhesion motifs; whilst gelatin used to control gelation and enhance cell adhesion. Based on our previous study ^26^, we use OA with a 50% degree of oxidation to achieve a suitable laminin-mimicking peptide concentrations in the prostate-specific alginate hydrogels. Two laminin-111 active peptides were selected for cellular attachment enhancement namely (i.e., IKVAV, AG73). IKVAV peptide, location residues α1 2097–2108, binds to α3β1 and α6β1 integrins, and known to promote cell adhesion, tumor growth, angiogenesis, metastasis, and protease activity ^12^. Laminin-111 peptide sequences are used to improve human prostate cancer cells (i.e., PC-3 cells, grade IV, adenocarcinoma) adhesion and human cancer-associated fibroblasts (i.e., CAFs), evaluating cell-material interaction by monitoring PC-3 phenotypes. PC-3 cells were selected as reported to highly express α 2, 3, 6, and β1 integrins^13,30^.

Based on clinical evidence on the role of stiffness in cancer progression ^9^ and our previous findings on breast cancer cells ^31,32^, we evaluated how stiffnesses impacted on PC-3 cells phenotypes (EMT markers and CD44 expression) and its correlation to their migratory and invasive phenotype, in presence or not of CAFs. Mechanical properties of hydrogels were controlled using different concentrations of crosslinker (CaCl_2_), with compression tests showing Young’s modulus in the range typical of PCa (i.e., 1-25 kPa), as reported in literature ^33–37^. Rheological tests confirmed printability of prostate-specific alginate formulations, and 3D *in vitro* models encapsulating PC-3 cells and CAFs were fabricated via EBB. PC-3 cells showed good viability and proliferation in the designed hydrogel up to 2 weeks, with significantly enhanced proliferation in softer hydrogel (i.e., E = 3.0 ± 0.5 kPa, hydrogel C1), in alginate functionalized with PEG and laminin peptides when compared to stiffer and not functionalized hydrogels (i.e., E = 12.6 ± 1.3 kPa, hydrogel A3) after 1 week. Expression of PCa progression markers (i.e., CD44, CD44v6, Vimentin, E-cadherin) was evaluated in PC-3 cells using 3D-printed models without/with CAFs, and showing dysregulated CD44 expression with increase of vimentin, linked to loss of CD44v isoforms and reduced E-cadherin in PC-3, indicative of invasive and metastatic traits.

This study presents a new technological platform that engineer the PCa TME both in physico-chemical and biological cues and aligns with recent guidelines of new approach methodologies providing more efficient ways to evaluate human cancer progression *in vitro*. Of note, the lack of PCa TME knowledge on biomechanical and biochemical composition limited the number of conditions herein presented. However, the manufacturing pipeline exploits technologies able to fabricate reproducible 3D *in vitro* models at a large scale offering a valuable platform to screen a large range of TME properties contributing to the generation of new knowledge better study of patient-specific PCa progression *in vitro*.

## 2. Materials and Methods

### 2.1 Materials

Sodium alginate (A_0_, 71238), sodium periodate (NaIO_4_, S1878), The 3-(trimethylsilyl)-2,2,3,3-tetradeuteropropionic acid (TMSP-d4, A14489.03), deuterium oxide (D2O, 7789-20-0), N-(2-Hydroxyethyl)piperazine-N′-(2-ethanesulfonic acid) (HEPES, H4034), gelatin type A (G, G1890), Pluronic®F127 solution (P2443), Nutrient Mixture F-12 Ham medium (N6658), Fetal Bovine Serum (FBS, F9665), 4% paraformaldehyde (PFA, 1004968350), L-glutamine (G751), Sodium bicarbonate (NaHCO3, S8761), Phosphate Buffer Solution (PBS, D1408), EMEM (M4655), Sodium pyruvate (P5280), Agarose (type 1, low EEO, A6013), Triton-X (648465), Saponin blocking buffer (47036), AccumaxTM (A7089), fibronectin (F2006) were all ordered from Sigma–Aldrich. 5,5’-Dithio-bis-(2-nitrobenzoic acid) (DTNB, 44889), cysteine hydrochloride monohydrate (44889), sodium chloride (NaCl, Fisher, 7647-14-5), calcium chloride (CaCl2, C/1400/53), sodium hydroxide (NaOH, 12963614), Live/Dead Kit (calcein AM and ethidium homodimer (EthD-1), L3224), Phalloidin Alexafluor488 (A12379), 4′-6-diamidino-2-phenylindole (DAPI, D1306) were all purchased from Thermo Fischer Scientific. Thiol-terminated IKVAV and AG73 peptides (custom-made by GenScript), poly-ethylene glycol (H_2_N-PEG-Mal, PBH-943, MW: 5 kDa, Creative PEGworks), Deep Blue Cell Viability™ Kit (424701, Biolegend), Cytopainter (ab138893, Abcam), Puromycin (A11138-03, Gibco), Cell dissociation buffer (13151014, Gibco) and collagen hydrogel precursor (50201, Ibidi).

### 2.2 Functionalization of alginate

#### 2.2.1 Preparation of Oxidized Alginate

Oxidized alginate (OA) was prepared following our protocol as reported in our previous studies ^26,27^. In this study, a 50% degree of oxidation (DO) was targeted; 8 g of sodium alginate was dissolved in 160 mL deionized water (dH_2_O) at RT overnight with mechanical stirring (400 rpm) using a tornado parallel reactor (RZR 2020, Heidolph, Germany), then 40 mL of 0.5 M NaIO_4_ (aq.) were gently added into the stirring alginate solution. The oxidation reaction was performed with continuous mechanical stirring (400 rpm) at RT for 6 h; then, the obtained OA_50_ solution (aq.) was purified by dialysis (Ultracel® 3kDa, PLBC07610) against dH_2_O at RT and in the dark to prevent further OA_50_ hydrolysis, changing the dH_2_O every day until completion of the purification process (i.e., dH_2_O conductivity < 8 μS/cm). OA_50_ solution (aq.) was freeze-dried (Chaist®, Alpha 2-4 LSC; 0.01 mbar, −80°C) until completely dry. The obtained OA_50_ powder was stored in the dark at RT and used within 18 months.

#### 2.2.2 Prostate-specific Alginate: Oxidized Alginate, PEG and Laminin-Mimicking Peptides

Prostate-mimicking alginates were functionalized considering integrins role in the initiation of PCa. As such, α-laminin thiol-terminated mimicking peptides were used in this study (i.e., IKVAV, AG73) The linear hetero-bifunctional PEG (H_2_N-PEG-Mal) was selected to link peptides to OA_50_ via maleimide-thiol Michael-type reaction ^38^. Thiol-terminated IKVAV was used as model peptide to assess alginate functionalization, then IKVAV and AG73 used for cell culture experiments. A 1:1 molar ratio of H_2_N-PEG-Mal and thiol-terminated IKVAV were allowed to react with constant stirring under continuous flow of argon gas (3 h, RT). The obtained product (H_2_N-PEG-IKVAV) was freeze-dried (0.01mbar, −80°C). The obtained PEG-peptide conjugate (H_2_N-PEG-IKVAV, H_2_N-PEG-AG73) was linked to OA_50_ via Schiff base reaction between the aldehyde groups (OA_50_) and the primary amino groups present on the PEG-peptide compound (**Figure 2A**). A 4% w/v OA_50_ solution in HBS (pH 8.5) was used varying the initial concentration of H_2_N-PEG-IKVAV (100-200 µM) with constant stirring (24 h, RT). The obtained solution was purified by dialysis (Spectra-Por^®^, MWCO 3.5-5 kDa, Z726273, Sigma–Aldrich, UK) against dH_2_O at RT and until conductivity < 8 μS/cm. The obtained functionalized alginate (OA_50_-PEG-IKVAV) was freeze-dried and stored until use.

### 2.3 Chemical characterization of prostate-specific alginates

#### 2.3.1 PEG-IKVAV Coupling Efficiency: Ellman’s Reagent

Ellman’s Reagent (DTNB) was used to quantify H_2_N-PEG-Mal and thiol-terminated IKVAV coupling efficiency. H_2_N-PEG-Mal (aq.) and IKVAV (aq.) were mixed and incubated at a 1:1 molar ratio (i.e., 500 µM) under Argon gas with constant stirring at RT for 90 min and 270 min. Samples were then mixed with DTNB following manufacturer’s instruction. Absorbance was measured at 412 nm using a plate reader spectrophotometer (BioTek, Synergy 2, NorthStar Scientific Ltd.). A standard calibration curve using cysteine hydrochloride monohydrate (aq.) in the molar range 0.25-1.5 mM to quantify unreacted peptide. The experiments were performed in triplicate (n=3) and for N=3 independent experiments.

#### 2.3.2 ^1^H NMR Spectroscopy

^1^H NMR was used to characterize alginate functionalization and to obtain the degree of functionalization (DF) using by Bruker Avance III-500 MHz NMR spectrometer, at 25 °C with 90° pulse proton angle, relaxation delay 2 seconds, and acquisition time 4.096 seconds, with total scans of 128 for each sample. A_0_ and OA_50_, and OA_50_-PEG-IKVAV samples were hydrated in 1 mL deuterium oxide (D_2_O) at a concentration of 10 mg/mL (2 min, 50°C). NH_2_-PEG-Mal, IKVAV, NH_2_-PEG-IKVAV were hydrated in 600 µL in D_2_O at a concentration of 5 mg/mL at RT. The DF was assessed using 4% OA50 solution (aq.) varying the concentration of NH_2_-PEG-Mal (i.e., 100 µM, 200 µM) allowing the reaction for 24h at RT. TMSP-d4 was used as an internal NMR standard to reference and quantify coupling efficiency. Prior acquisition, all samples were freeze-dried (0.01 mbar, −80°C) until completely dry, then re-dissolved in D_2_O. Spectra were analyzed by topspin software (Bruker, 4.1.0).

### 2.4 Preparation of prostate-specific Alginate Hydrogels

Alginate and gelatine solutions (aq.) were dissolved in HBS at RT and 37°C respectively, at concentrations reported in **Table 1**. Alginate and gelatin solutions were sterile filtered using a 0.22 µm PES filter (Millex®GP) and a 0.45 µm PVDF filter (Millex®-HV), respectively. Sterile solutions were gently mixed, paying attention not to generate air bubbles and obtaining *biomaterial ink **A*** (A_0_-gelatin-OA_50_) and *biomaterial ink **B***, (A_0_-gelatin-OA_50_-PEG-IKVAV/AG73). CaCl_2_ solutions (aq.) were prepared at concentration of 0.1 and 0.3 M, and sterile filtered using a 0.22 µm PES filter prior use. For mechanical testing, cylindrical hydrogels were prepared by adding 550 µL of A_0_-OA_50_-G solution (aq.) in a cylindrical mold (i.e., 8 mm diameter, 8 mm height) allowing gelatin sol-gel transition and Schiff’s base formation (1 h, 4°C) fixing the shape; the physical crosslink step was performed incubating each sample in 3 mL of CaCl_2_ solution (aq.) allowing ionic gelation (10 min, 37°C). CaCl_2_ solution (aq.) was removed, and samples washed with HBS solution (3x, 10 min) to remove any excess of CaCl_2_. Prior testing, hydrogels were incubated in F-12 medium at 37°C in a humidified atmosphere with 5% CO_2_ for 24 h for complete swelling and equilibration ^27^. Of note, the molds were incubated with a sterile 3% (w/w) Pluronic®F127 solution (aq.) for 10 min and left air dry overnight in a sterile condition to form a homogeneous coating; before use, the bottom of the mold was sealed with PCR film (LightCycler® 480 sealing foil, USA). For cell culture tests, samples were incubated in CaCl_2_ solutions (aq.) allowing physical gelation (10 min, 37°C). CaCl_2_ solution (aq.) was removed, and the obtained hydrogel samples washed with HBS solution (3 times, 5 min, RT), to ensure removal of CaCl_2_.

**Table 1.** Composition of prostate-specific alginate precursor solutions and hydrogels. For hydrogel preparation, precursors solutions with unmodified alginate (A_0_), oxidized alginate (OA_50_), modified alginate (OA_50_-PEG-peptide) and gelatin (G) were mixed and crosslinked with CaCl_2_ solutions (aq.) allowing gelation at 37°C, 10 min.

| Sample ID | *A*<sub>0</sub> (% w/v) | *OA*<sub>50</sub> (% w/v) | *OA*<sub>50</sub>-PEG-pept (% w/v) |  | *G* (% w/v) | CaCl<sub>2</sub> (μM) |
| --- | --- | --- | --- | --- | --- | --- |
|  |  |  | *IKVAV* | *AG73* |  |  |
| **A** | 1 | 1 | 0 | 0 | 3 | 0 |
| **A1** | 1 | 1 | 0 | 0 | 3 | 100 |
| **A3** | 1 | 1 | 0 | 0 | 3 | 300 |
| **B** | 1 | 0 | 1 | 0 | 3 | 0 |
| **B1** | 1 | 0 | 1 | 0 | 3 | 100 |
| **B3** | 1 | 0 | 1 | 0 | 3 | 300 |
| **C** | 1 | 0 | 0.5 | 0.5 | 3 | 0 |
| **C1** | 1 | 0 | 0.5 | 0.5 | 3 | 100 |
| **C3** | 1 | 0 | 0.5 | 0.5 | 3 | 300 |

Concentrations reported in table are the final concentrations in each formulation.

### 2.5 Mechanical and physical properties

#### 2.5.1 Rheological tests

Rheological tests were first performed to measure flow properties and gelation kinetics of *biomaterial inks* A and B (**Table 1**) prior gelation (i.e. hydrogel precursor solutions) using the Thermo Scientific^TM^ HAAKE^TM^ MARS rheometer. Values were recorded by HAAKE RheoWin Job Manager (Version 4.87.0002). Flow properties were evaluated using 20 mm cone plate geometry (C20/1° Ti L, 222-1877, Thermo Scientific), with a gap (plate-to-plate) adjusted to 0.1 mm, at shear rate range 0.1 – 300 s^-1^. All tests were performed at 37 °C.

A 35 mm parallel plate geometry was used to measure the gelation kinetics, the gap (plate-to-plate) was set to 0.1 mm prior measurements. Oscillatory frequency and strain were kept constant at 10% and 1 Hz, respectively. Storage modulus (G′) and loss modulus (G′′) were measured at 37 °C to measure the gelation kinetics and identify the gelation time of *biomaterial inks*.

Additional rheological tests were performed to measure the viscoelastic properties of the produced hydrogels (**Table 1**) by carrying out oscillation amplitude strain sweeps in the range 0.01-100% strain at constant frequency of 1 Hz using serrated 8 mm parallel plate geometry, at 37 °C and with the gap (plate-to-plate) adjusted to the height of each sample (4-5 mm). Retrieved data were analyzed using DIN 51810-2 recommended settings in RheoWin Data Manager (version 4.87.0002) to determine the values of G’ and G’’ in the linear viscoelastic range (LVR). All tests were performed using n=3 samples, for N=3 independent experiments.

#### 2.5.2 Compression Tests

Compression tests were performed using cylindrical samples (d= 8 mm, h= 5 mm) to measure the compressive moduli (i.e., Young’s modulus, E) of hydrogels, and performed as previously reported by our group ^31^. Prior to each test, the diameter of each sample was measured using a caliper and further used to calculate the surface. Compression test was performed using the Texture Analyzer (Stable Micro Systems Texture Analyzer, TA-XT plus) equipped with 0.5 N load cell (Stable Micro Systems, Load: 538855). Samples were placed on the Texture Analyzer stage and compressed with 0.1 mm/s probe speed without pre-compression. The selected compression speed guarantees to work within the linear viscoelastic region of soft hydrogels. For each sample, E was derived from the stress-strain curve as the slope of the curve within the range of 0-5 % strain. All tests were performed using n=3 samples, for N=3 independent experiments.

#### 2.5.3 Porosity

Porosity of alginate-based hydrogels was investigated using the solvent replacement method ^24^. Hydrogels were prepared as cylinders, freeze-dried (0.01mbar, −80°C) and weighted (w_1_). Samples were immersed in excess of absolute ethanol (24 h, RT), and weighed after excess ethanol on the gel was removed by blotting (w_2_). Porosity (%) was calculated using **Equation 1**, in which *ρ* is the density of absolute ethanol and *V* corresponds to the volume of the sample. All tests were performed using n=3 samples, for N=3 independent experiments.

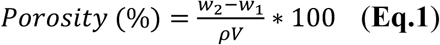

### 2.6 Prostate-specific 3D *in vitro* models

#### 2.6.1 Cell Culture

PC-3 human prostate cancer cells were purchased from American Type Culture Collection (ATCC^®^ CRL-1435^™^) and cultured in complete F-12 Ham cell culture medium (7% (v/v) FBS, 2 mM L-glutamine). PC-3 cells were discarded upon reaching passage 80. hTERT PF179T CAF human prostate cancer associated fibroblast cells were purchased from American Type Culture Collection (ATCC® CRL-3290TM) and maintained in complete EMEM (contain Earle’s Balanced Salt Solution, non-essential amino acids and supplemented with 10% (v/v) FBS, 2 mM L-glutamine, 1 mM sodium pyruvate, 1500 mg/L sodium bicarbonate 7.5% and 10 mg/mL puromycin. hTERT PF179T CAFs (from now on referred to simply as CAFs) were discarded upon reaching passage 25. Co-culture experiments were performed using complete F-12:EMEM medium (1:1 volume ratio) as reported in (**Supplementary Information, SI.5**). All cells were routinely cultured in the incubator (37°C, 5% CO_2_).

#### 2.6.2 Preparation of prostate bioinks and stromal bioinks

Sterile *biomaterial ink **A*** (A_0_-gelatin-OA_50_) and *biomaterial ink **C***, (A_0_-gelatin-OA_50_-PEG-IKVAV/AG73) were used to prepare bioinks. *Prostate* and *stromal bioinks* were obtained by homogeneously mixing biomaterial inks with PC-3 and hTERT PF179T CAF at a concentration of 1×10^6^ cells/mL in a 1:1 ratio, respectively.

#### 2.6.3 Microbeads: Prostate-specific and stromal-specific 3D *in vitro* models

*Prostate-specific* and *stroma-specific bioinks* were transferred in a 1 mL syringe and extruded through a 27G nozzle dropwise in the CaCl_2_ crosslinking solution allowing gelation (10 min, RT). After crosslinking, formed hydrogel beads were collected using a cell strainer (CSS-010-040, Biofil, UK), transferred to HBS solution and incubated (10 min, RT) to remove any excess of CaCl_2_. Finally, cell encapsulating hydrogel beads (i.e., *prostate-specific microbeads*, *stroma-specific microbeads*) were transferred to complete cell culture media and cultured in incubator (37°C, 5% CO_2_).

#### 2.6.4 Suspension bath-based Bioprinting: Engineered 3D prostate 3D *in vitro* models

Agarose fluid gels were used to support the printed construct during printing process before crosslinking with CaCl_2_ solutions ^24,39^. Briefly, 0.5% w/v agarose fluid gels were prepared in dH_2_O by cooling autoclaved agarose form 100 °C to 25 °C under a constant shear of 700 rpm for at least 6 hours. In a typical experiment, 2 mL/well of sterile fluid agarose gels were then loaded into 6-well tissue culture plates and 3 mL *bio/material ink* loaded into a printing cartridge using a piston to avoid undesired leakages and maintain sterility. The 3D Discovery Evolution bioprinter (RegenHU, Switzerland) was used to print engineered 3D prostate-specific *in vitro* models. First, printing parameters were optimized using *biomaterial ink* using conditions specified in **Table SI.1** and selecting optimal ones. After printing, the printed constructs were incubated (10 min, RT) allowing Shiff’s base reaction (**Supplementary Information SI.3**), then agarose gel was gently removed leaving a very thin layer to prevent any possible damage. CaCl_2_ crosslinking solution (aq.) was added to the well, allowing ionic crosslinking (10 min, 37°C, 5% CO_2_). Finally, constructs were gently washed with HBS (3x, 10 min).

For printing of engineered 3D prostate-specific *in vitro* models, 3 mL of *Prostate bioink* or *Stromal bioink* were loaded into a printing cartridge, and the printing performed with the same setting previously described and with optimized printing parameters.

### 2.7 Characterization of engineered prostate-specific 3D *in vitro* models

#### 2.7.1 Cell Metabolic Activity

Deep Blue Cell Viability™ assay following manufacturer’s instruction and at selected time points (i.e., 1, 4, and 7 days). Briefly, one *prostate-specific microbead* was places in a 96 well plate, the cell culture media gently removed and replaced with 200 µL of 10% v/v deep blue viability solution in complete cell culture media, samples were incubated (2 h, 37°C, 5% CO_2_), and then 100 µL of media was transferred to new 96 well plate. Samples were read immediately using a plate reader spectrophotometer (BioTek, Synergy 2, NorthStar Scientific Ltd.) at an excitation range of 530-570 nm and emission range of 590-620 nm. The measurements were performed using n=5 samples, for N=3 independent experiments.

#### 2.7.2 Cells viability: Live/dead assay

Live/Dead assay following manufacturer’s instruction in *prostate-specific microbeads* and *engineered 3D prostate in vitro models*. At the selected time point, encapsulated cells in hydrogels were incubated (1 h, 37°C, 5% CO_2_), washed twice with HBS, fixed with 4% v/v PFA (30 min, RT), and followed by final HBS washing. Images were acquired using confocal fluorescence microscope using (Ex/Em 488/525 nm) and (Ex/Em 570/600 nm) filters to detect calcein (live cells, green) and ethylene homodimer (dead cells, red), respectively. Z-stack images were post-processed using CQ1 software.

#### 2.7.3 Marker expression: Flow cytometry

Expression of selected markers (i.e., CD44, CD44v6, E-cadherin and vimentin) was performed using the flow cytometer (BD Fortessa X-20) at day 3 and 7. PC-3 cells cultured on standard tissue-culture plates were used as control (i.e., 2D). Briefly, 2D controls were washed with PBS, incubated with cell dissociation buffer (10 min, 37°C, 5% CO_2_) ^40^; whereas for 3D *in vitro* models, cells were recovered by dissolution of hydrogels as previously reported by Rosa *et al*. ^41^ retrieved cell aggregates were disrupted with AccumaxTM solution (10 min, RT). All cells were centrifuged at 600 g (5 min, RT), gently resuspended in blocking buffer (5% (v/v) FBS in PBS) and incubated for 30 minutes on ice. For intracellular markers detection (i.e., vimentin), cells were fixed with 4% PFA (10 min, RT), washed thrice with PBS, followed by a permeabilization step with 0.1% w/v Saponin in blocking buffer (30 min, RT). Cells were then incubated on ice with primary antibody (45 min) and then with the secondary antibody (45 min) at concentrations reported in **Table 2**. Dead cells exclusion was performed by DAPI staining (i.e., incubation of cells with 1 μg/mL DAPI in PBS) and positive cells analyzed using FlowJo software (v10.8.0, BD). Single live cells gating was performed, obtaining the number of positive cells for each marker and calculating the median fluorescence intensity (MFI). Median intensity data are reported as an average of N=3 independent biological experiments.

**Table 2.** Used markers and corresponding dilutions of isotype control, primary antibodies and secondary antibodies.

| Marker | Isotype control | Primary antibody | Secondary antibody |
| --- | --- | --- | --- |
| CD44 | APC Rat IgG2b, $\kappa$<br>Isotype Ctrl (400611,<br>BioLegend) | APC anti-mouse/human<br>CD44 Antibody (103012,<br>BioLegend) | N/A |
|  | (dilution 1:800) | Dilution (1:800) |  |
| CD44v6 | Mouse IgG1 Negative<br>Control antibody<br>(MCA928, Bio-Rad) | Mouse anti Human<br>CD44v6 (MCA1730,<br>Bio-Rad) | BV786 Rat Anti- Mouse<br>IgG1, (742480, BD) |
|  | (dilution 1:10) | (dilution 1:100) | (dilution 1:150) |
| Vimentin | Rat IgG2A APC Isotype<br>control (IC006A, R&D<br>systems) | APC Rat anti-human<br>Vimentin (IC2105A,<br>R&D systems) | N/A |
|  | (dilution 1:10) | (dilution 1:20) |  |
| E-cadherin | APC anti-human CD324<br>(B 263116, BioLegend) | APC Mouse IgG1<br>isotype (400121,<br>BioLegend) | N/A |
|  | (dilution 1:30) | (dilution 1:30) |  |

N/A = not applicable

#### 2.7.4 Cell Aggregates: Size and Shape quantification

At selected time points (i.e., day 7, day 14), PC-3 cells in *prostate-specific microbeads* were fixed with 4% PFA solution (aq.) (15 min, RT), washed trice with 1× HBS, permeabilized with 0.1% Triton-X solution in HBS (15 min, RT), washed trice with PBS and incubated with a DAPI (1 μg/mL) and phalloidin-AlexaFluor488 (1:80 v/v dilution in HBS) solution (45 min, RT). *Prostate-specific microbeads* were washed thrice with HBS and stored in HBS with antimycotic-antibiotic for image acquisition using (Ex/Em 405/447 nm) and (Ex/Em 488/525 nm) filters to detect DAPI (nuclei, blue) and phalloidin 488 (F-Actin, green), respectively. Aggregates were identified thresholding and converting images to a binary format; then selected using the ‘create selection’ function and finally the area and shape characteristics of the selected aggregates were measured using the ‘Analyze Particles’ plug-in in ImageJ (v1.53a). Circularity was used as a shape descriptor to assess the resemblance of each object to a perfect circle, with a value of 1.0 indicating a perfect circle and 0.0 indicating a highly elongated shape.

#### 2.7.5 Cell Adhesion assay

Collagen and fibronectin were used to coat 8-well chambers slides (80826, Ibidi) according to manufacturer’s protocol. Briefly, collagen type I was diluted in 17.5 mM acetic acid solution (aq.) to 35 μg/mL, 200 μL added to each well and incubated (1 h, RT) allowing absorption. Similarly, 200 μL/well of 20 μg/mL fibronectin solution (aq.) was used and incubated (1 h, RT). After this incubation step, solutions were removed, wells washed with sterile PBS and left to air dry in sterile conditions (1h, RT). PC-3 cells pre-conditioned in *prostate-specific microbeads* (7 days) were recovered by dissolving the hydrogel network as reported in ^41^ and seeded on uncoated (control), collagen or fibronectin coated surfaces at a density of 1×10^4^ cells/cm^2^ and allowed to adhere to surfaces (1 h, 37°C, 5% CO_2_). Cells were washed twice with PBS to remove not-adhered cells, fixed with 4% PFA (15 min, RT), washed thrice with PBS, and further permeabilized with 0.1% Triton-X in PBS (15 min, RT). Cells were incubated with 200 μL/well of a 1 μg/mL DAPI in PBS and a Phalloidin (1:80 ratio of dilution) (45 min, RT), washed trice with PBS and stored in HBS with antimycotic-antibiotic for image acquisition using (Ex/Em 405/447 nm) and (Ex/Em 488/525 nm) filters to detect DAPI (nuclei, blue) and phalloidin 488 (F-Actin, green), respectively.

### 2.8 Image Acquisition

#### 2.8.1 Brightfield

Brightfield images of encapsulated cells in hydrogels beads were acquired using a fluorescent inverted microscope (Leica DMI6000, Leica Microsystems, UK), connected to a 5.5 Neo sCMOS camera (Andor, UK). For image capturing µManager software (v.1.46, Vale Lab, UCSF, USA) was used. For acquisitions, a dry 2× objective (PLAN 2.5/0.07, Leica), dry 10× objective (PL 10/0.3 PH1, Leica), and a dry 20× objective (PL 20/0.5 PH2, Leica), were used.

#### 2.8.2 Immunofluorescence

Images were acquired using the confocal microscope (CQ1 Confocal Imaging Cytometer Yokogawa) coupled with sCMOS camera (2000×2000pixel, 13.0×13.0mm). Using microlens-enhanced wide-view Nipkow disk. For acquisitions, 4×,10×, and 20× objectives and (Ex/Em 405/447 nm), (Ex/Em 488/525 nm) and (Ex/Em 570/600 nm) filters were used. For all samples Z-stacks were acquired with a 5 μm z-step.

### 2.9 Statistical Analysis

All results were reported as mean ± SD. For statistical analysis all results were analyzed with two-way ANOVA using GraphPad Prism v9.5.1. P-values were set at four different significance levels: (*p ≤ 0.05, **p ≤ 0.01, ***p ≤ 0.001, ****p ≤ 0.0001).

## 3. Results and Discussion

### 3.1 Biomaterial ink and prostate-specific hydrogels characterization

#### 3.1.1 Oxidized Alginate physico-chemical properties

Oxidation of alginate was confirmed by ^1^H NMR (**Figure 1**) spectra. ^1^H NMR spectra of OA_50_ show the typical alginate fingerprint with peaks in the region from 3.6 to 4.0 ppm, corresponding to the protons of both G- and M-units. As the oxidation involves only the G units, OA_50_ spectrum shows a signal at 4.2 ppm with distinctive peaks at 5.35 and 5.60 ppm (**Figure 1**), both attributed to hemiacetalic proton formed after oxidation ^42^. Aldehyde signals expected at 9.7 ppm ^43^, was not detected possibly due to the hemiacetal formation of aldehyde groups with the adjacent hydroxyl groups as explained previously ^26^.

**Figure 1.**
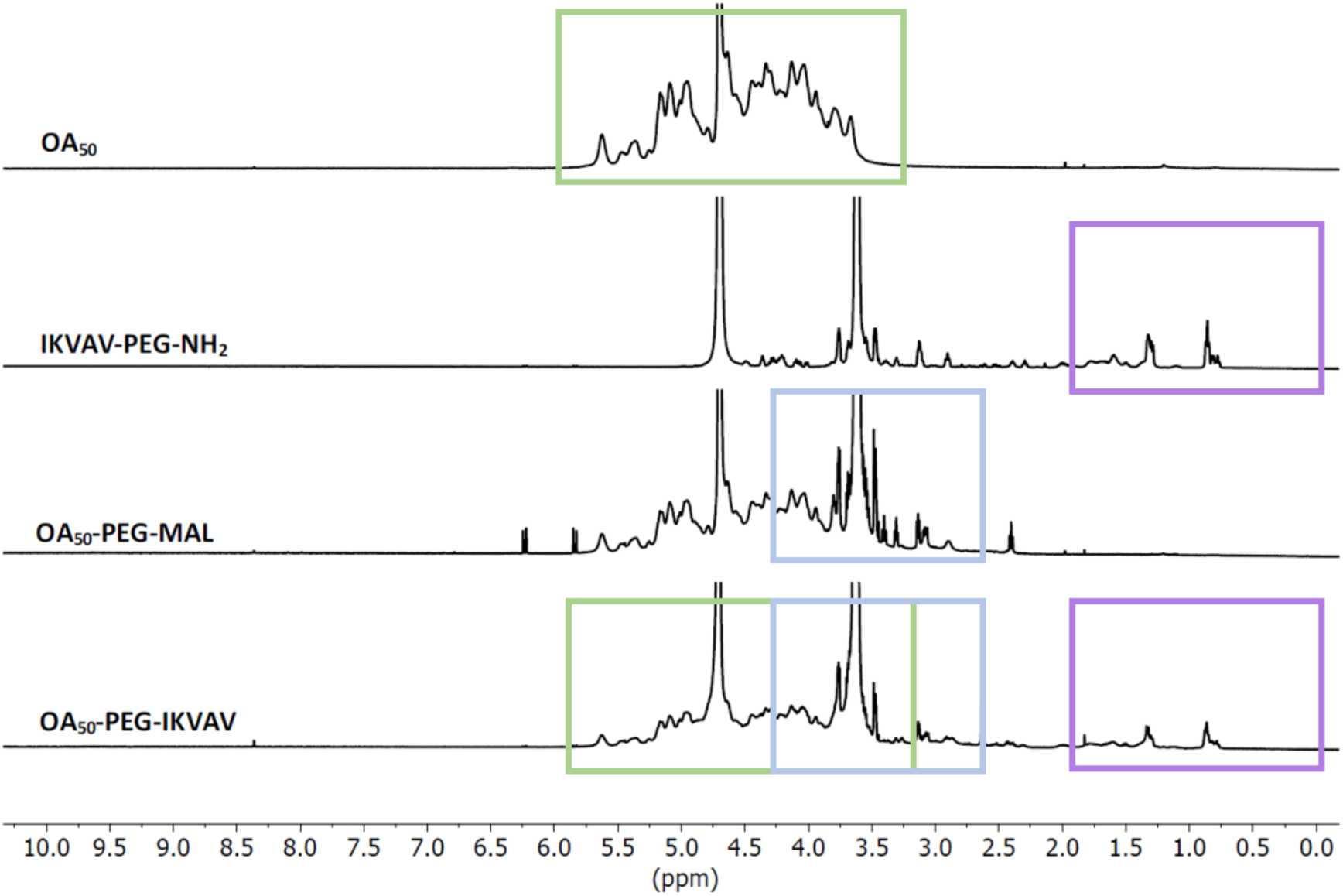
Characterization of functionalized alginate. ^1^H-NMR spectra of oxidized alginate OA_50_, IKVAV-PEG-NH_2_, OA_50_-PEG-MAL and OA_50_-PEG-IKVAV.

#### 3.1.2 Functionalized Alginate: Coupling Oxidized Alginate-PEG-Peptide

The functionalization of OA_50_ with the selected peptide (i.e. IKVAV) was assessed via H^1^-NMR spectra (**Figure 1**), whilst the quantification of the DF was done by Ellman’s assay. H^1^-NMR spectra were used to confirm the coupling and synthesis of OA_50_-PEG-IKVAV (**Figure 1**). Bi-functional H_2_N-PEG-Mal and thiol-terminated IKVAV coupling was assessed. Prior, a verification of the PEG-peptide coupling (IKVAV-PEG-NH_2_) was confirmed (**Figure 1**): H_2_N-PEG-Mal spectrum (**Figure SI.1A**) shows characteristic peaks of PEG (3.0-4-0 ppm), and around 6.70 ppm representing maleimide group ^44^, with peak at 5.98 ppm and 6.38 ppm attributed to double bond protons of the maleimide group ^45^. IKVAV spectra reports characteristic peaks of valine, isoleucine and alanine in the range 1.5-0.5 ppm ^46^ (**Figure SI.1A**). Of note, the IKVAV-PEG-NH_2_ intermediate product shows clearly all the peaks corresponding to both IKVAV and PEG (**Figure 1**), thus confirming the coupling of IKVAV to the PEG. Further, the reduced signal assigned to maleimide group (at δ 5.9–6.5 ppm) indicates that a reaction with thiols occur (Michael-type addition reaction). **Figure 1** finally confirms coupling of IKVAV-PEG-NH_2_ with OA_50_: the region >2.0 ppm in the OA_50_-PEG-IKVAV matches the IKVAV signal, which is not displayed in the corresponding OA_50_-PEG-Mal signal (used as control).

To quantify the conjugation between H_2_N-PEG-Mal and OA_50_, the effect of H_2_N-PEG-Mal concentration was assessed by comparing the ratio of the integrals for H^1^-NMR reference (TMSP) protons to the methyl protons from PEG’s terminal methoxy group (δ ∼3.2ppm): a coupling efficiency of approx. 70% was obtained for both concentration (**Figure SI.1B**), used to set the concentration of H_2_N-PEG-IKVAV (or any other coupled peptide to H_2_N-PEG-Mal) to be used for the conjugation with OA_50_ and quantified to 100 µM, confirming what reported in literature ^38^. The coupling efficiency of selected IKVAV peptides with PEG was quantitated by Ellman’s Reagent, using cystine for the standard calibration curve. This assay was used to monitor the reaction kinetics, and sampling reaction products after 1.5 h and 2.5 h. It was observed an increase in the coupling efficiency from 80% (reaction time of 1.5 h) to 99% after 2.5 h reaction (*data not shown*). Based on these results, it was decided to incubate the peptide with PEG for 3 hours to complete the conjugation by Michael-type addition reaction to ensure the complete coupling between the reagents.

### 3.2 Physical characterization of Prostate specific Alginate-based Hydrogels

#### 3.2.1 Porosity and pore size

Porosity of hydrogels is a physical property that influence prompt water uptake and cell migration ^47^. Hydrogel porosity was evaluated by solvent replacement (**Figure 2A**). Hydrogels **A1** and **B1** have higher porosity (approx. 10%), reflecting the lowest crosslinking density linked to the lowest CaCl_2_ (aq.) concentration, whereas hydrogels **A3** and **B3** have lower porosity (**Figure 2A**). As expected, it is possible to evidence the role of crosslinker concentration with the lowest (i.e., 100 μM CaCl_2_) returning the highest porosity. of note, hydrogel **A3** is the sample with the lowest porosity: this value is linked to the higher concentration of crosslinker (300 μM CaCl_2_) and the presence of additional crosslinks between gelatin and OA_50_ (**Supplementary Information SI.3, Figure SI.2**).

**Figure 2.**
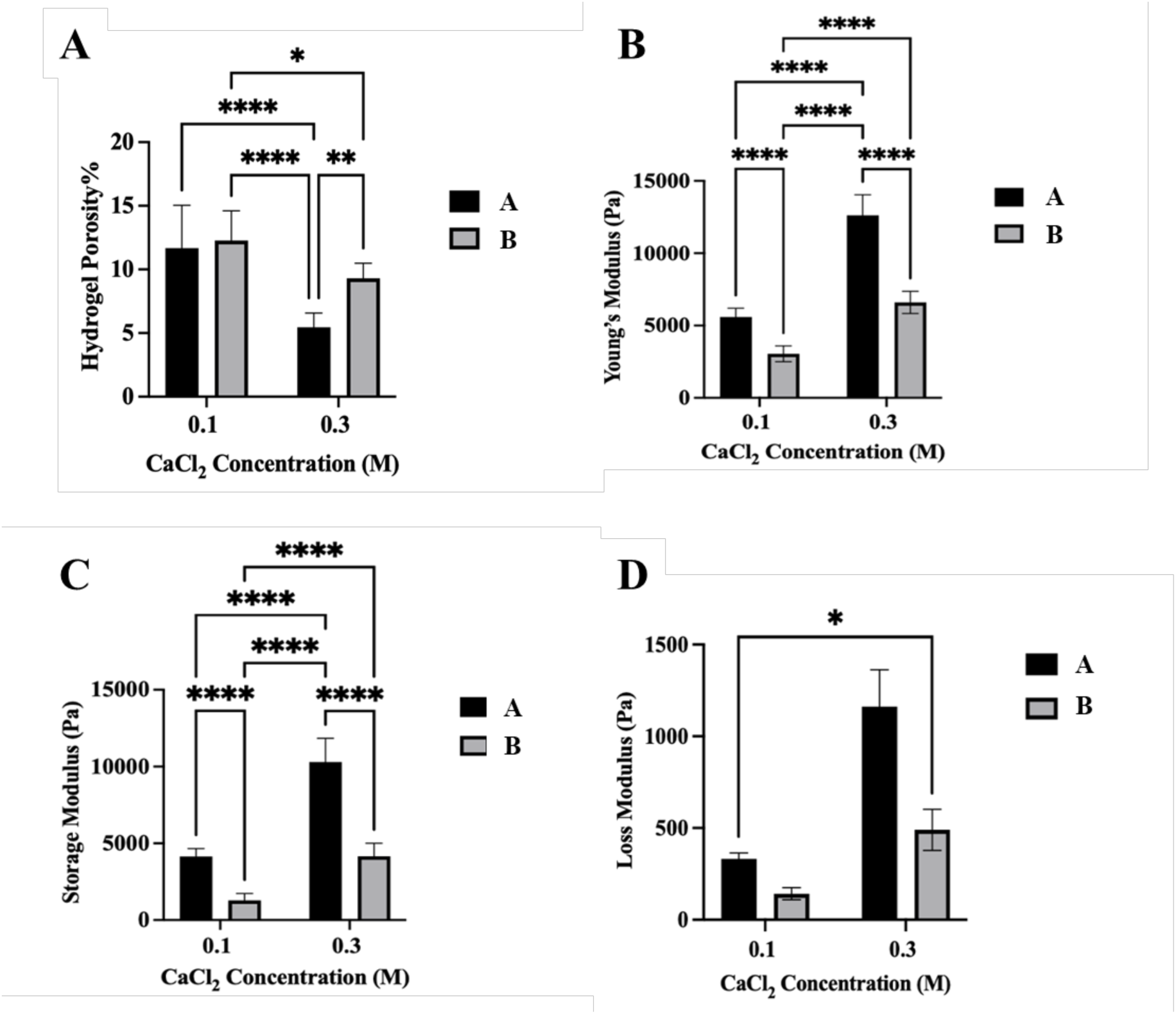
Physical and mechanical properties of alginate-based hydrogels. **A)** Porosity percentage of alginate-based hydrogels. **B)** Young’s modulus measured by uniaxial compressive tests. All hydrogels and CaCl_2_ concentration are significantly different (p ≤ 0.0001). **C)** Storage and **D)** loss modulus measured with Rheometer at constant frequency (1 Hz). All hydrogels and CaCl_2_ concentration are significantly different (p ≤ 0.0001). Data presented as mean ± SD (N=3, n=3) for all samples. P-values represented as *p ≤ 0.05, **p ≤ 0.01, ***p ≤ 0.001, ****p ≤ 0.0001.

#### 3.2.2 Mechanical Properties of Prostate-specific Hydrogels

Tissue stiffening in PCa is typically linked to variations of ECM composition, and mainly due to increased collagen deposition, controlling mechano-signaling and transduction to cells during tumor progression ^6,35^. Based on values reported in literature on the mechanical properties of human prostate tissues ^33–37,48^, we formulated hydrogels addressing both healthy prostate tissue and advanced/metastatic PCa.

Compression tests were performed with no initial pre-stress and to quantitate the Young’s modulus on the different hydrogels (**Table 1**) with measured values in the range of 2.5 – 13.0 kPa (**Figure 2B**). As expected, results show that E is proportional to CaCl_2_ (aq.) concentration, hence with crosslinking density. Moreover, the presence of PEG on alginate chains causes a decrease in measured E, possibly due to the increased water content in hydrogels. As anticipated, hydrogels **A1** and **A3** have increased crosslinked density due to the covalent crosslinks OA_50_ and gelatin (Schiff’s base formation between amino groups and aldehyde groups, **Supplementary Information SI.3, Figure SI.2**) that increases E when compared to hydrogel B1 and B3. In fact, hydrogel **A1** is stiffer than hydrogel **B1** (5.6 ± 0.6 kPa vs 3.0 ± 0.5 kPa, P ≤ 0.0001) even if crosslinked with the same CaCl_2_ (aq.) concentration (100 μM); similar trend is observed between hydrogels **A3** and **B3** (12.6 ± 1.3 kPa vs 6.6 ± 0.7 kPa, P ≤ 0.0001) when using a higher CaCl_2_ (aq.) concentration (300 μM).

Rheological tests results align with the trend shown in compression tests and with measured storage moduli (G′, **Figure 2C**) higher in hydrogels **A1** and **A3** (4.1 ± 0.5 kPa, 10.3 ± 1.5 kPa) when compared to hydrogels **B1** and **B3** (1.3 ± 0.4 kPa, 4.2 ± 0.8 kPa), with increased G’ at higher CaCl_2_ (aq.) concentrations (P ≤ 0.0001) and reduced G’ in the presence of PEG in alginate chains (P ≤ 0.0001). The loss modulus (G”, Figure 2D) measured in shear sweep tests is following the same trend as per G’, with the highest value in hydrogel A3 (1.2 ± 0.2 kPa) and the lowest in hydrogel B1 (0.1 ± 0.0 kPa). CaCl_2_ (aq.) concentration is again proportional to the loss modulus: hydrogels A1 and A3 (0.3 ± 0.0 kPa vs 1. ± 0.2 kPa, P ≤ 0.0001) have higher loss modulus than hydrogel B1 and B3 (0.1± 0.0 kPa vs 0.5 ± 0.1 kPa, P ≤ 0.0001).

Of note and as reported, the addition of PEG to alginate chains causes hydrogels higher loss modulus, this is being reported to be proportional with PEG concentration and molecular weight ^49^, which was not investigated in this study.

### 3.3 Manufacturing of Prostate-specific 3D *in vitro* models

#### 3.3.1 *Biomaterial Inks*: rheological properties, printability and gelation kinetics

Biomaterial inks should meet specific requirements in the flow properties (i.e., shear thinning behavior), allowing extrudability and shape retention for ease manufacturing of 3D models with bioprinters ^50^. Here, flow properties of prostate *biomaterial inks* were evaluated with rheological test, measuring the dynamic viscosity in response to shear rate and assessing their shear thinning properties ^51^. Two *biomaterial inks were* tested as precursors of hydrogel **A1** and **A3** (i.e., *biomaterial ink **A***) and hydrogels **B1** and **B3** (i.e., *biomaterial ink **B***), both showing shear thinning behavior (**Figure 3A-B**).

**Figure 3.**
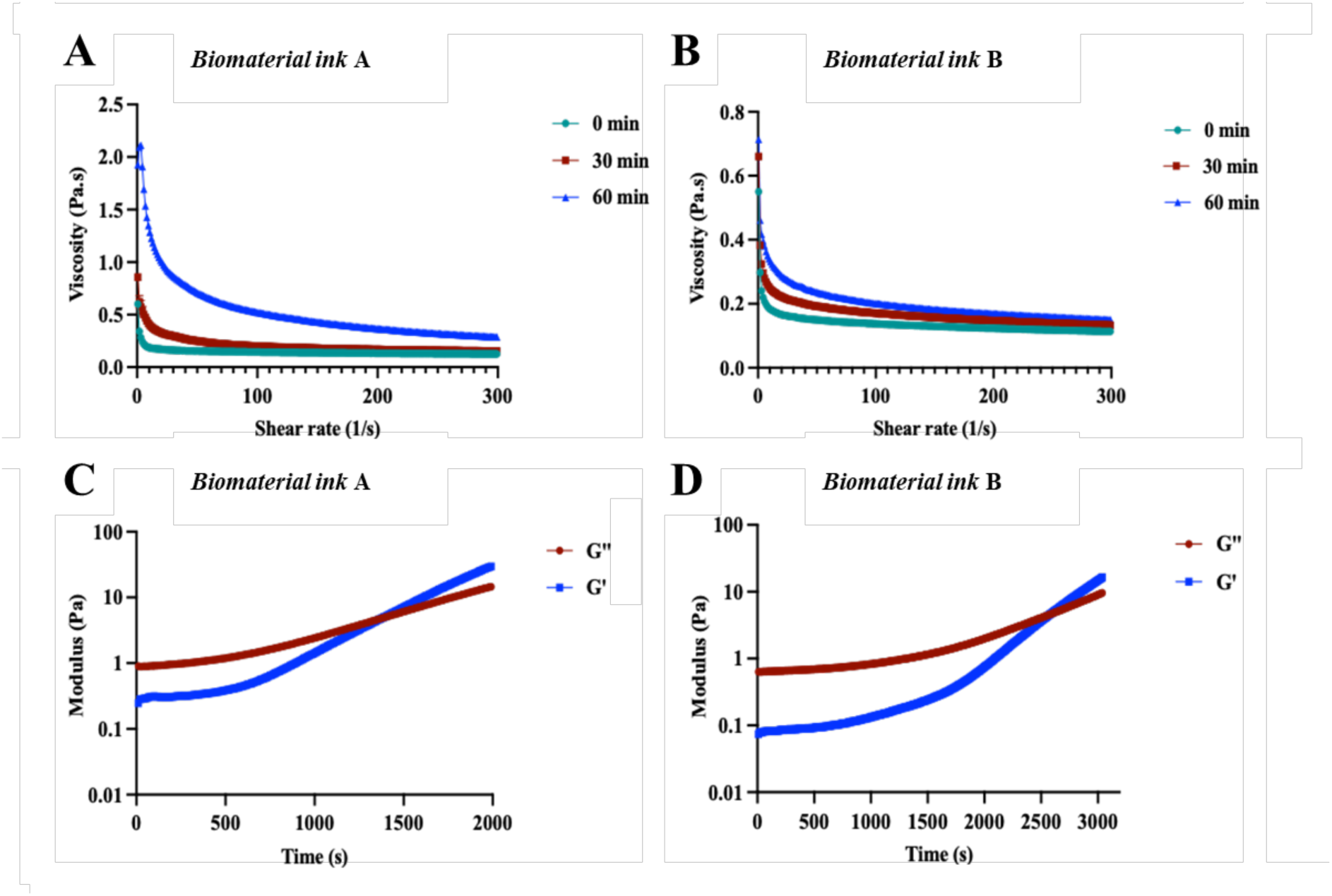
Flow Curve and Gelation Kinetics. **A-B)** Viscosity as a function of shear rate (37 °C) for biomaterial inks A and B**; C-D)** Time sweep rheology of biomaterial inks. Over time storage modulus (G′) and loss modulus (G′′) were measured at 37 °C to obtain gelation time post mixing the gel components where gelation time was recorded at crossover of G′ and G′′. Data are presented as mean of (N=3, n=3).

As per hydrogels formulation design, *biomaterial inks* have two components reacting to form covalent crosslinks (Shiff’s base reaction between OA_50_ and gelatin) and allowing hydrogel formation: this reaction causes variation of *biomaterial ink* viscosity over time. Therefore, gelation kinetics was monitored though rheological tests and to identify the printability window, in which no effect of the Shiff’s base reaction is observed. The gelation kinetic of prostate biomaterial *inks* show G’ values of *biomaterial ink **A*** higher than *biomaterial ink **B*** already after mixing (0.25 ± 0.16 and 0.07 ± 0.03 respectively). Rheological tests show an average gelation time *biomaterial ink **A*** of 24.5 ± 3.5 min at the crossing point (i.e., G’ = G”: 5.9 ± 0.9 Pa, **Figure 3C-D**). *Biomaterial ink **B*** has a gelation time of 43.4 ± 2.1 min at the crossing point (i.e., G’ = G”: 4.7 ± 0.8 Pa, **Figure 3C-D**). Such difference in gelation time and crosslinking point value (5.9 ± 0.9 Pa vs 4.7 ± 0.8 Pa) confirms the lower availability of aldehyde groups to react with primary amines of gelatin. As a result, and to ensure constant dynamic viscosity values during printing, *biomaterial inks* are prepared and used within 10 min after preparation.

#### 3.3.2 Prostate-specific 3D *in vitro* models: Microbeads

Cell viability and re-organization of PC-3 cells and CAFs was evaluated in prostate-specific alginate hydrogel microbeads, assessing any possible influence of biomechanical and biochemical elements of the TME. Representative brightfield images of *prostate-specific microbeads* prepared with *prostate bioinks* crosslinked as hydrogel **C3** are shown at different timepoints (i.e., day(s) 1, 4, 7) in **Figure 4A**. Regardless the *bioink* used (either *prostate* or *stroma bioink*), microbeads show uniform diameter (approx. 1 mm) stable over time (**Supplementary Information SI.6, Figure SI.5**). Viability of PC-3 cells is confirmed by Live/Dead assay (**Supplementary Information SI.6, Figure SI.6**). PC-3 metabolic activity increases over time (day(s) 1, 4, and 7, **Figure 6B**), with higher proliferation PC-3 cells rate in hydrogel **C1**. PC-3 cells aggregate differently when cultures in *prostate-specific microbeads* varying composition and mechanical properties are shown in **Figure 4C**, as previously observed in our previous studies in breast-specific microbeads ^31^; cell aggregates vary in size and shape over time (day 7, and day 14), showing a *in vivo-like* tumor structure as response and adaptation to the microenvironment ^52^.

**Figure 4.**
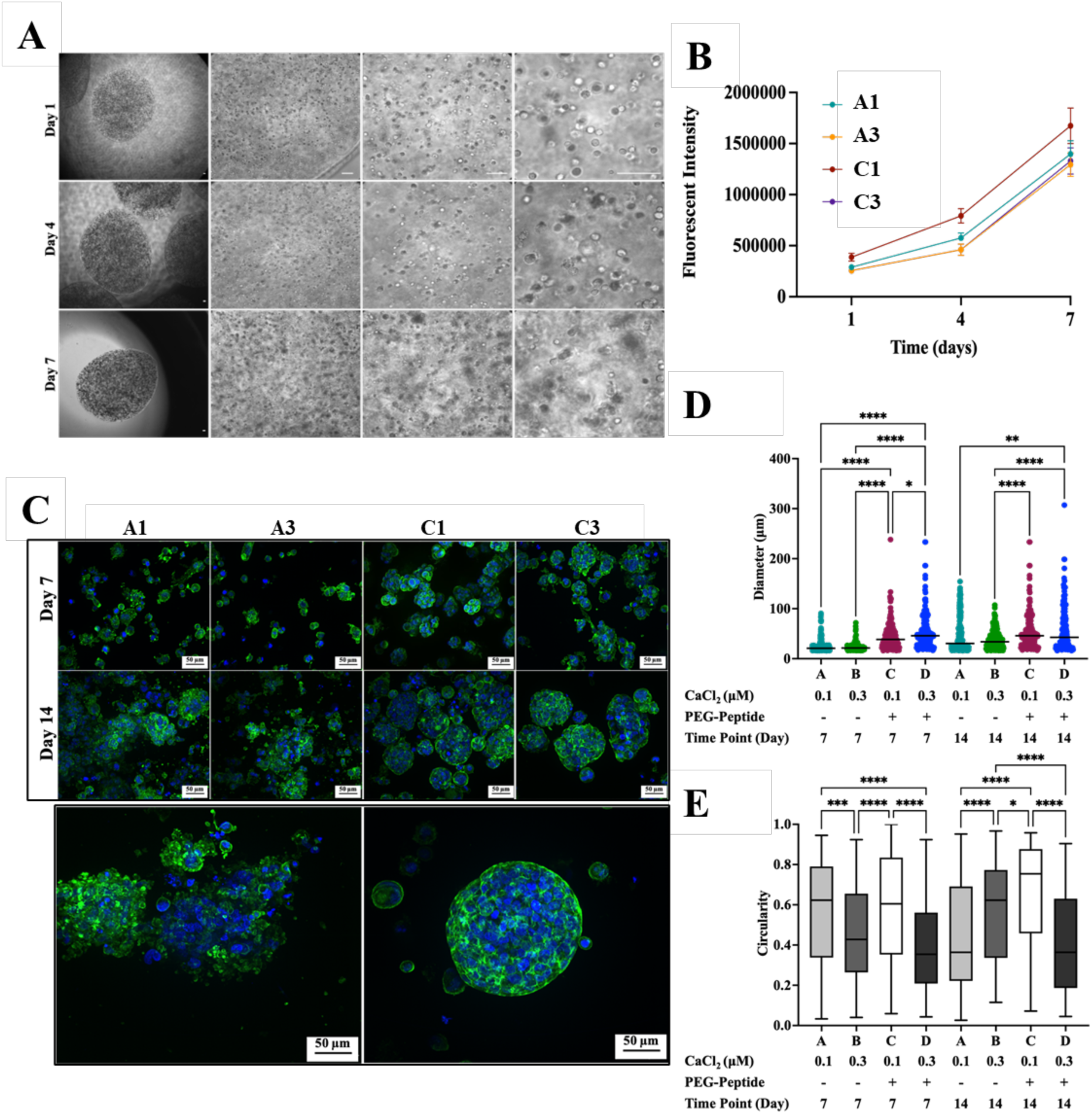
PC-3 prostate 3D in vitro models. **A**) Representative images of PC-3 cells encapsulated in hydrogel C3. Images acquired with a 2×, 10×, 20×, and 40× objective respectively, at day1, 4, and 7. Scale bars 100 μm. **B)** Cell viability was measured by deep blue fluorescence at different time points (day 1, 4, and 7). Statistical analysis at day 7 using Two-way ANOVA returned (P ≤ 0.0001) for (hydrogel 3) vs (hydrogel 2), and (hydrogel 3) vs (hydrogel 1). Data are presented as mean ± SD (N=3, n=3), P-values represented as *p ≤ 0.05, **p ≤ 0.01, ***p ≤ 0.001, ****p ≤ 0.0001. **C)** Immunofluorescent images of PC-3 cells stained with DAPI (nuclei, blue) and Phalloidin (F-Actin, green) showing cell aggregates. Below Representative images of PC-3 cell aggregates with low (left) and high (right) circularity values. Scale bars: 50 μm. **D)** Quantification of diameter (μm) of aggregates. E) Quantification of circularity of aggregates. The labels in graphs are denoted as 2D (2D TCP plate), A hydrogel A1, B hydrogel A3, C hydrogel C1, D hydrogel C3. Values are represented as mean and SD, at least n=200 aggregate for each tested condition. P-values represented as *p ≤ 0.05, **p ≤ 0.01, ***p ≤ 0.001, ****p ≤ 0.0001.

As shown in **Figure 4C**, PC-3 cells started to form cellular aggregates at day 7 in all hydrogels with pronounced formation in hydrogels **C1** and **C3** when compared to hydrogels **A1** and **A3** (p ≤ 0.0001). In the latter hydrogels, few cell aggregates are observed, with majority of cells remain dispersed in the hydrogel or form smaller aggregate. In all hydrogels, cells form larger aggregates over time (day 7 vs day 14), suggesting a sustained cell proliferation. Shape of aggregates is scored as circularity, for which 1 is assigned to objects perfect fitting a circular structure and 0 to irregular ones. PC-3 cells form grape like structures with high circularity in hydrogel **C1** (p ≤ 0.0001 vs hydrogel **C3**), and as also observed in other works using chitosan-chondroitin sulphate and Matrigel scaffolds ^53,54^. A reduced circularity is observed at day 14 in both hydrogel **C1** and **C3** when compared to day 7 (**Figure 4D-E**), which suggests a morphological transformation of round aggregates into a more invasive structures at day 14 and observed bridge-like structures connecting cell aggregates ^54^.

#### 3.3.3 Bioprinting Engineered 3D prostate *in vitro* models: suspension bath-based bioprinting

Maintaining a good shape fidelity and shape retention while using low-viscosity biomaterial inks is challenging: to tackle this problem suspension bath-based bioprinting was used to print low viscosity *biomaterial inks* ^55^. Briefly, suspension bath-based bioprinting relies on a fluid-like gel with homogeneous particle distribution, shear-thinning, and self-healing properties to enable the movement of the printing needle without disrupting the printed construct until crosslinked. The printing parameters were optimized for *biomaterial ink **A*** and *biomaterial ink **B***, without inclusion of cells (*bioinks*). *Biomaterial inks* were printed at different conditions (**Table SI.1**), and filament diameter calculated from images (**Figure 5A-B**). A continuous filament of *biomaterial ink* **A** is obtained with low extrusion pressure (20 to 30 kPa) and high feed rate (7.5 to 10 mm/s) (**Figure 5B**). As expected, and for both *biomaterial inks*, at all extrusion pressures increased feed rate led to a decrease in filament diameter and reaching dimensions comparable to the nozzle diameters in few conditions (**Figure 5C-D**), as also reported elsewhere ^50^.

**Figure 5.**
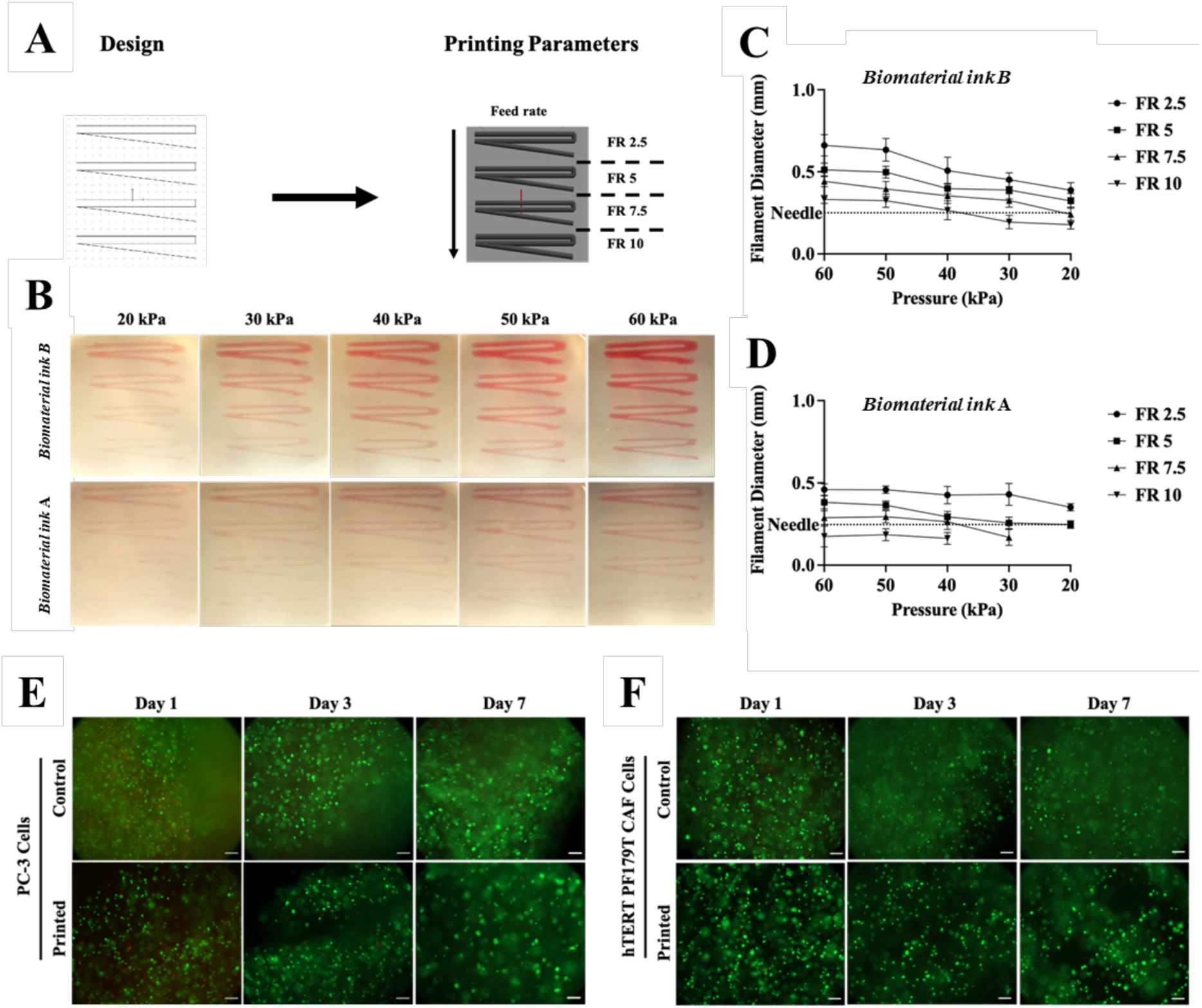
3D printed prostate specific in vitro models. **A-B) 3D bioprinting parameters optimization via extrusion-based bioprinting.** **C-D)** Quantification of filament diameter. Data are plotted as mean ± SD (N=2, n=3). **E-F) Live/dead assay of printed cells.** Printed PC-3 and CAF cells at day 1 and 7 were compared to control encapsulated cells to evaluate the effect of shear stresses during the printing process. In the image live proliferative cells are displayed in green, whereas dead cells in red. Scale bars: 200 μm.

hTERT PF179T CAFs were co-cultured with PC-3 cells to better recapitulate the prostate TME and evaluate the role of stromal cells in PCa progression. To assess the feasibility in controlling the spatial location of PCa cells (i.e., PC-3) and CAFs (i.e., hTERT PF179T CAFs), *prostate bioinks* and *stromal bioinks* were printed using optimal printing parameters selected as 20 kPa extrusion pressure, and 7.5 mm/s feed rate. Cells viability assessed using Live/Dead staining 1 day and 7 days after printing confirms cell viability of PC3 cells higher than 90% (**Figure 5E-F**) and confirms the possibility to use bioprinting to further engineer 3D prostate *in vitro* models.

### 3.4 Effect of prostate-specific TME on PC-3 prostate cancer cells phenotypes

Novel alternative methods (NAMs) is form of 3D engineered prostate-specific *in vitro* models where herein used to assess their possibility to link with clinical data and possibly predict clinical outcomes. PC-3 cells showed different behaviors when cultured in different prostate-specific microbeads (**Figure 4**), forming larger and more circular cell aggregates when cultured in prostate specific hydrogels (i.e., **C1**, **C3**). Engineered PCa TMEs were obtained using suspension bath-based bioprinting and controlling the architecture, the biochemical composition and mechanical properties of prostate ECM, as well as localization and density of PCa and stromal cells (i.e., PC3 and CAFs were printed with a 1:1 cell number ratio). The impact of PCa cells and the prostate ECM on metastatic potential was evaluated on PC-3 cells behaviors (e.g., adhesion, migration, invasiveness) and phenotypes at different time points and allowing PC-3 cells adaptation to the engineered TME.

#### 3.4.1 Adhesion on different ECM-mimicking interfaces

The first key process of PC-3 adaptation to different prostate-mimicking ECM (i.e., hydrogels **A1**, **A3**, **C1**, **C3**) and presence of stromal cells (i.e., monoculture, co-culture with CAFs) evaluated was PC-3 cells ability to adhere to different surfaces coated with ECM relevant components, namely collagen and fibronectin, and compared to uncoated surfaces. Quantitative evaluation of PC-3 cells adhesion was performed comparing number of adhered cells and their spreading area among selected surfaces and after 1 week PC-3 cells adaptation to the engineered TMEs. This 2D tests were designed to validate cellular adhesion which normally anticipates migratory and invasive behavior.

PC-3 cells were allowed to adhere for 1 hour to different surfaces (i.e., tissue culture treated, collagen, fibronectin) after preconditioning in different engineered microenvironments, then stained to evaluate their morphology (**Figure 6A**). Among all the engineered prostate 3D *in vitro* models, the highest number of adhered PC-3 cells is recorded after preconditioning in hydrogel **C3** (E = 6.6 ± 0.73 kPa), where the presence of CAFs in all surfaces tested did not significantly impact on the number of adhered PC-3 cells (**Figure 6B**) and with the highest number of adhered cells identified on collagen coated surfaces (**Figure 6C**). Of note, conditioning TME does not highly impact on PC-3 cells adhered to uncoated wells, whereas a certain level of correlation between conditioning TME and adhesion surface is observed with fibronectin and collagen coated surfaces (hydrogel **C3** > **A3** ≥ **C1** > **A1**); similar trends are observed in both coated surfaces in presence of CAFs. Overall, it is not just the biomechanical properties (e.g., stiffer ECM) that drives cell adhesion capacity, but rather the biochemical elements (i.e., presence of α-laminin peptides) in which PC-3 cells are preconditioned. These results align with formation of PC-3 cell aggregates (**Figure 4C**) with less circular and more invasive morphologies in hydrogels **A3** and **C3**, suggesting that a stiffer ECM promotes expression of an invasive cells phenotype. Cell spread area (**Figure 6D**), further confirm that both hydrogel stiffness (E ≤ 6 kPa) and composition (presence of α-laminin peptides) increase the migration and invasion potential of PC-3 cells. Of note, PC-3 cells protrusions formation are observed as indicative of preliminary stages of migration and invasion (*data not shown*)^48,56^.

**Figure 6.**
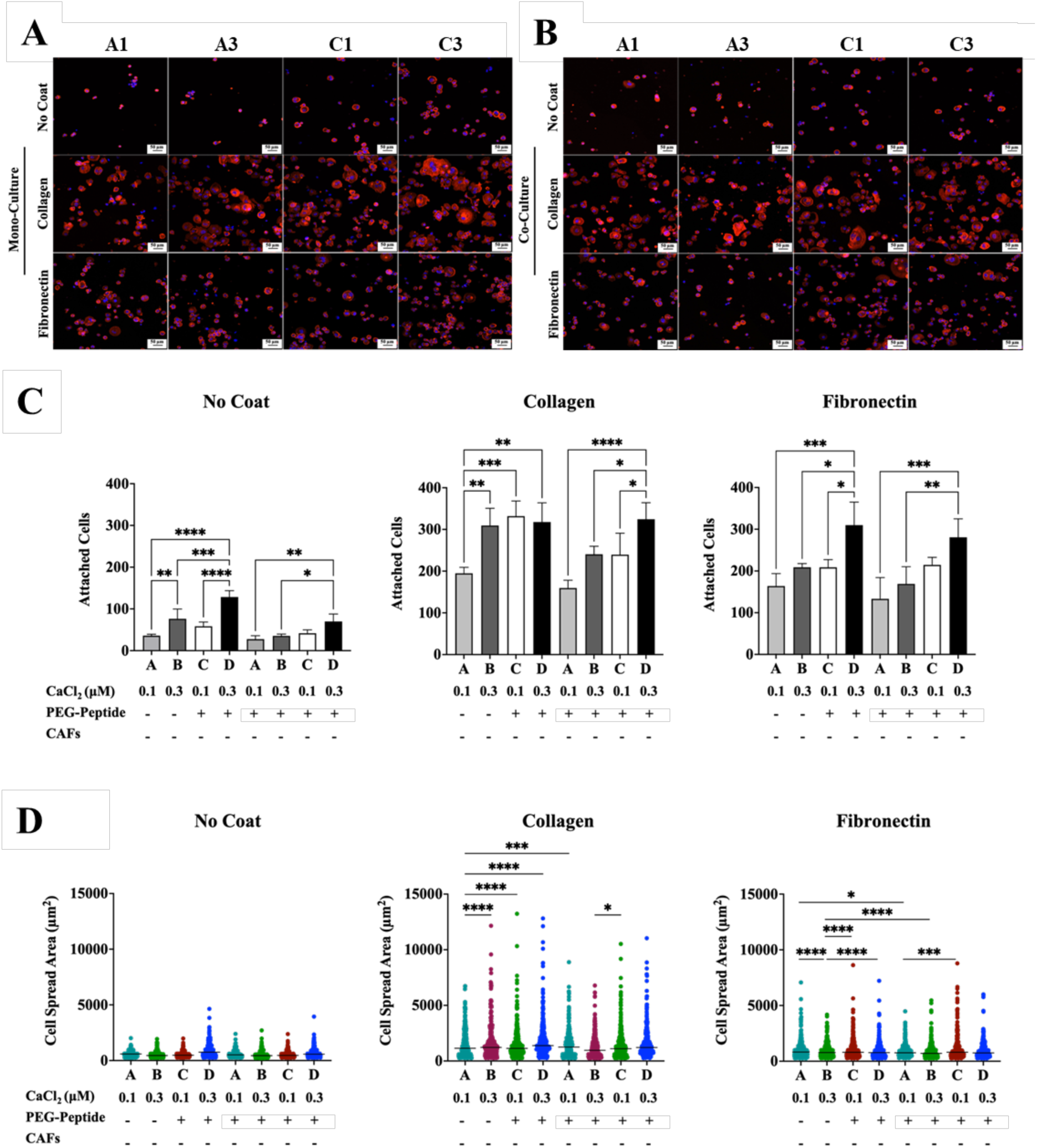
PC-3 cell adhesion on different substrates without/with CAFs. **A-B)** Immunofluorescent images of PC-3 cells preconditioned in hydrogels without/with co-culture with CAFs cells. Cells are stained with DAPI (nuclei, blue) and Phalloidin (F-Actin, red) then let to adhere to different surfaces, non-coated, collagen-coated or fibronectin-coated surfaces (Scale bars: 50 μm). **C)** Count of attached cells to different surfaces in different models. **D)** Dot-plot representations of individual cell spread area and mean in different surfaces when cells allowed to adhere for 1 h. The labels in graphs are denoted as A (hydrogel A1), B (hydrogel A3), C (hydrogel C1), D (hydrogel C3). P-values represented as *p ≤ 0.05, **p ≤ 0.01, ***p ≤ 0.001, ****p ≤ 0.0001.

#### 3.4.2 Epithelial-To-Mesenchymal Transition Markers and CD44 Expression

Examining phenotypic changes in PCa cells is essential to understand aggressiveness and metastatic potential; with EMT being recognized as pivotal mechanism in PCa progression and predictive in determining clinical outcomes. Among the EMT markers used to evaluate tumor progression in PC-3 cells, we selected to evaluate the expression of E-cadherin, Vimentin, CD44s and CD44v6 ^57,58^.

Clinical evidence suggests a correlation between specific TME traits and PCa aggressiveness, with a shared underlying thread being the higher Gleason score the more aggressive the cancer. In this, ECM interaction with PCa cells drives their phenotypic variations over time, as well as the presence of stromal cells ^59,60^. We found that PC-3 cells express increased levels of Vimentin when co-cultured with CAFs, with a limited impact of the ECM traits (**Figure 7A**). Of note, PC-3 cells are found positive for Vimentin (>80%) in all tested conditions and time points and in line with what reported by Xu et al. ^61^. Of note, PCa cells positive for CD44 express higher level of vimentin ^57^, when PCa cells interact with CAFs tend to express higher levels of vimentin: this corroborates with expression of CD44 of PC-3 in this study (**Figure 7C**). We observed, as reported in literature, that the Epithelial-To-Mesenchymal Transition (EMT) in PC-3 cells is also linked to loss of CD44v isoforms, which coincide with reduced E-cadherin (**Figure 7B**) and increased mesenchymal markers like vimentin ^57^. In all tested conditions, E-cadherin MFI is found constant in PC-3 cells, with higher number of positive cells found in presence of CAFs (approx. 40% vs approx. 50%) ^62^.

**Figure 7.**
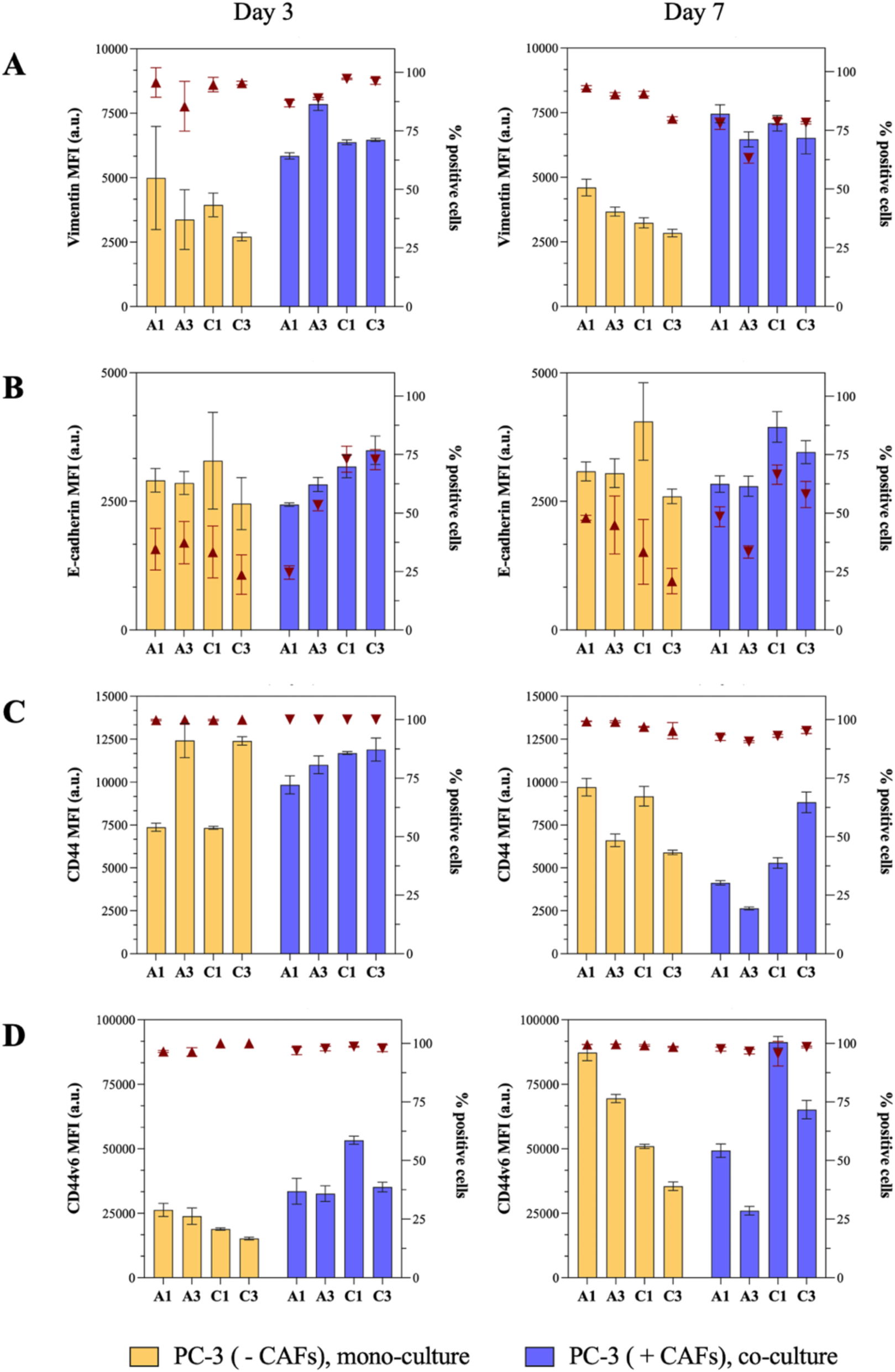
Flow cytometry analysis of EMT markers and CD44. Flow cytometry quantification in PC-3 cells encapsulated in different hydrogel formulas of **(A)** median fluorescence intensity (MFI) of Vimentin, **(B)** MFI of E-cadherin, **(C)** MFI of CD44, **(D)** MFI CD44 v6. The percentage of positive cells (%) is represented in each plot (dark red triangles). Values are represented as mean and SD of N=3 independent experiments.

Dysregulated CD44 expression characterizes most human cancers, including PCa. We observed an increase in CD44 expression in stiffer hydrogel **A3** (p ≤ 0.0001) and hydrogel **C3** (p ≤ 0.0001) with E value ranging from 12.6 to 6.6 kPa after 3 days in engineered TME when PC-3 are cultured alone and compared to softer hydrogels hydrogel **A1** and **C1** with E value ranging from 5.6 to 3.0 kPa (**Figure 7C**). At the same time point, and when PC-3 cells are co-cultured with CAFs, this trend is not observed with high CD44 expression and with >95% found positive for the standard CD44 isoform. Interestingly, the trend is inverted after 7 days for PC-3 in mono-culture: in softer ECM (i.e., hydrogels **A1**, **C1**) CD44 expression increases and higher than PC-3 cells cultured in stiffer ECM (i.e., hydrogels **A3**, **C3**). An overall reduction of CD44 expression in PC-3 co-cultured with CAFs is observed, with no correlation with biomechanical and biochemical traits of hydrogels.

Clinically, CD44 expression tends to decrease as PCa progresses to higher Gleason grades and more poorly differentiated, aggressive tumors. Such association suggests that with higher vimentin and lower E-cadherin expression (i.e., PC-3 EMT), reduced CD44 expression is associated with aggressive traits in PC-3 cells. Overall, in presence of CAFs, CD44 decreases in PC-3 cells showing a more invasive/aggressive phenotype after 7 days, suggesting a higher metastatic potential of PC-3 cells. Further investigation correlating the CD44/CD24 and PC-3 cancer stemness may be of use to better interpret their invasiveness. On this note and based on our previous findings on breast cancer cells invasiveness ^31,32^, we investigated the expression of CD44v6 isoform typically implicated in tumor cells invasion and metastasis ^57^. Expression of CD44v6 in PC-3 cells (> 95%, across all the conditions tested) confirms the aberrant increase of CD44 variant isoforms with the loss of CD44s expression as PCa progresses. Markers of tumor differentiation and progression show that among the different TMEs tested stiffer and laminin-enriched ECMs (i.e., hydrogel **C3**) is overall associated with invasive traits of PC-3 cells, with and without CAFs. This aligns with adhesion results, in which stiffer and laminin-enriched ECM promotes higher PC-3 adhesion on substrates (**Figure 6**). To follow up on this, we further investigated migration and invasiveness of PC-3 cells observing similar behaviors: at day 3, PC-3 cells from hydrogel **C3** have significantly higher migration capability compared to other TMEs (**Figure SI.7**, **Figure SI.8**) and a possible influence of CAFs on the migratory ability of PC-3 cells. We assessed PC-3 cells ability to invade collagen ECM at early time points (**Supplementary Information SI.8, Figure SI.8**), but none of the condition tested at this early time point evidence a distinguished invasive trait of PC-3 cells. To better investigate the role of TME on cells invasive phenotypes, longer observational time points may be required.

## 4. Conclusions

This work shows that engineered prostate 3D *in vitro* models are useful technologies to study PCa cells’ phenotype in alginate tumor-like hydrogels controlled in biomechanical and biochemical properties. 3D bioprinting can be exploited to manufacture specific 3D *in vitro* prostate tissue hence studying how the TME directs PCa cells aggressiveness and invasiveness. Alginate was modified to include laminin-like domains (i.e., AG73, IKVAV) and physical hydrogels formed with controlled viscoelastic properties (2.5-13 kPa, matching healthy prostate or tumoral tissues) to evaluate expression of tumor markers (i.e., CD44, Cd44v6, vimentin, E-cadherin) and metastatic potential of PC-3 cells.

Described 3D PCa models displayed good viability and proliferation of PC-3 cells up to 7 days, with PC-3 cells cultured in hydrogels enriched in laminin showing better adhesion and invasiveness. We found that ECM properties, mainly stiffness, impact on PC-3 phenotypes than the presence of CAFs; however, prolonged co-culture with stromal cells may enhance PCa cells’ metastatic potential.

Overall, the proposed models are suitable for large scale and high-throughput studies, enabling ease testing of EMT and other clinically relevant markers to be linked with clinical data, hence predict clinical outcomes. Refinement of engineered prostate 3D *in vitro* models will more closely mimic the human TME in controlling biomechanical, biochemical and cells distribution (tumor, stromal) assessing metastatic progression or response to chemotherapeutics towards personalized medicine.

