## Supplementary Information for "Bio-fabricated alginate tumor-like hydrogels to enhance understanding of prostate-specific micro-environments *in vitro*"

### SI.1 Chemical characterization of modified alginate: $^{1}$H-NMR

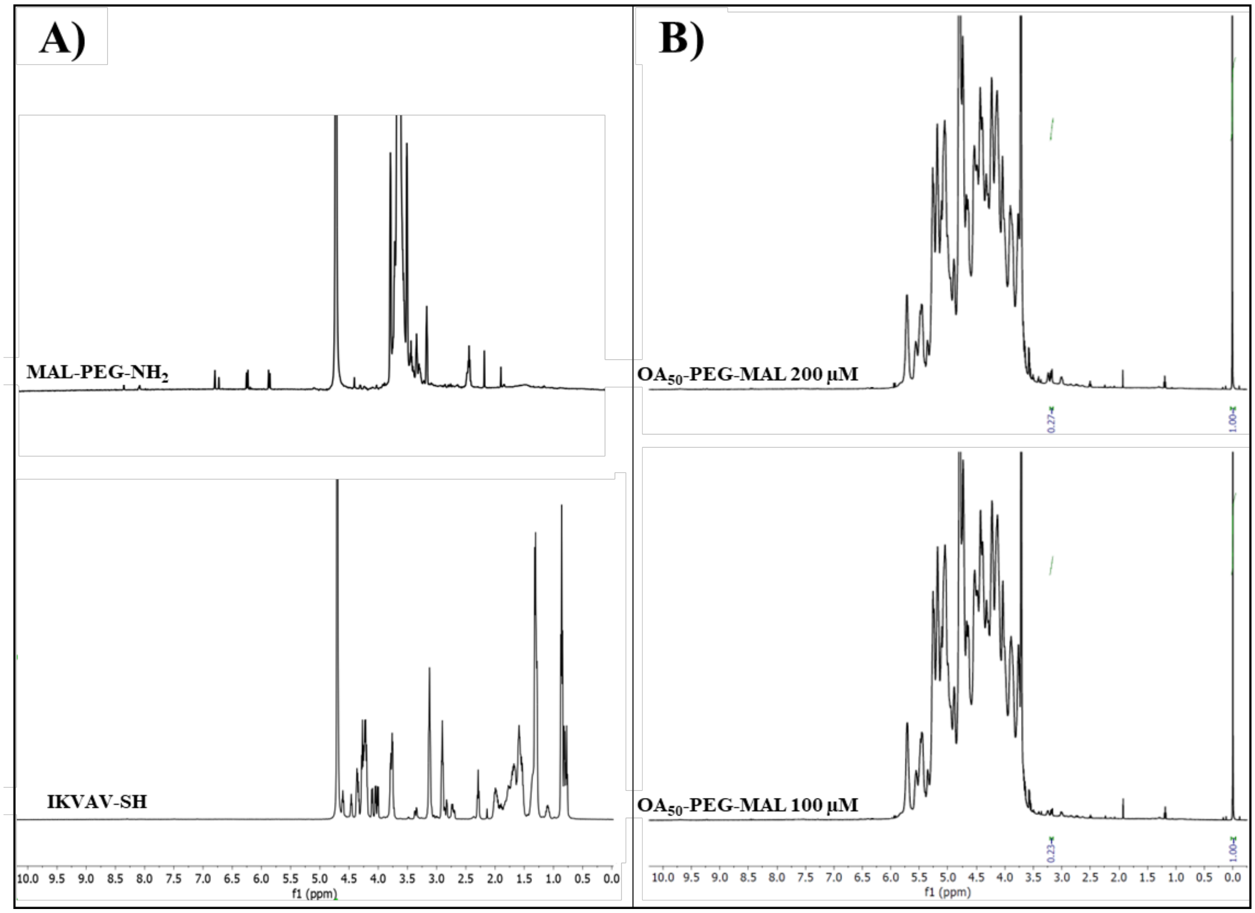


**Figure SI.1.** $^{1}$H-NMR spectra of **A)** IKVAV-SH and MAL-PEG-$NH_2$ and **B)** $OA_{50}$-PEG-MAL.

### SI.2 Alginate Oxidation

Following our previous study, sodium periodate was used to oxidize sodium alginate and form aldehyde groups on alginate backbone, obtaining the oxidized alginate (OA) with known degree of oxidation (DO). In this study a 50% DO was targeted to ensure functionalization with a high density of laminin-mimicking peptides. Of note, oxidized alginate and its degree of oxidation are abbreviated as $OA_{DO}$, whilst unmodified alginate is abbreviated as $A_0$. Obtained $OA_{50}$ has a molecular weight of 22.2 ± 0.6 kDa and an aldehyde concentration of approx. 6.5 μM/mg, and confirming values reported in our previous study [1].

### SI.3 Conjugation between Oxidized Alginate ($OA_{50}$) and Gelatin (G)

Aldehydes groups on $OA_{50}$ were quantified at a concentration of 6.5 μM/mg [1], and used to react with available ε-amino groups of lysine or hydroxylysine in gelatin (Schiff's base reaction). Gelatin FT-IR spectrum shows the characteristic bands of gelatin at 1626 $cm^{-1}$ attributed to C=O stretching of amide I, and 1520 $cm^{-1}$ of N-H deformation for amide II. After mixing with alginate-based solutions (i.e. $A_0$ /$OA_{50}$, $A_0$/$OA_{50}$-PEG-IKVAV), biomaterial inks are obtained. FT-IR spectra of biomaterial ink **A** and biomaterial ink **B** show the presence of the corresponding peaks of Schiff's base at around 1614-1617 $cm^{-1}$ and 1536-1546 $cm^{-1}$. Schiff's base absorption band in biomaterial ink **A** is more pronounced probably due to the availability of more aldehyde groups to crosslink with ε-amino groups of gelatin. The broader Schiff's base band at 1617 $cm^{-1}$ in biomaterial ink **B** is possibly obtained because of overlapping with the band at 1626 $cm^{-1}$of amide I of un-crosslinked gelatin. Furthermore, the characteristic peak of gelatin at 1520 $cm^{-1}$ representing amide II was not detected in both hydrogel formulas verifying the involvement of this group in the crosslinking reaction.

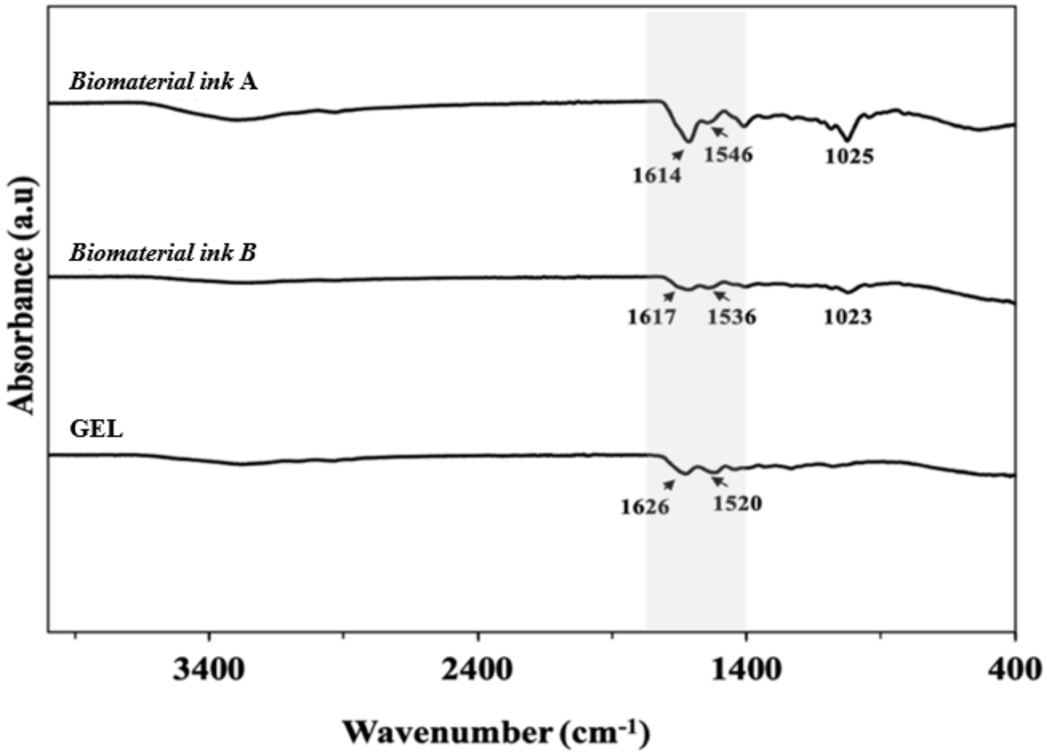


**Figure SI.2. Oxidized Alginate and Gelatin Conjugation characterized with FT-IR spectra in the range of wavenumber 400-4000 cm$^{-1}$**. FT-IR spectrum for gelatin (GEL), biomaterial ink A, and biomaterial ink B. Characteristic bands of gelatin were at 1626 cm$^{-1}$ attributed to C=O stretching of amide I, and 1520 cm$^{-1}$ of N-H deformation for amide II.

### SI.4 Manufacturing of prostate-specific hydrogels

**Table SI.1**. Printing parameters used with 3D Discovery Evolution bioprinter to optimize manufacturing of prostate-specific 3D models. The process was performed at constant temperature (22 ± 3°C).

| **Printing parameters** | **Selected values** |
|---|---|
| Nozzle type | 25G (Steel, cylindrical) |
| Printing speed (mm/s) | 2.5; 5.0; 7.5; 10.0 |
| Extrusion pressure (kPa) | 20, 30, 40, 50, 60 |

**Table SI.2. Alginate-based hydrogels composition and mechanical properties.** Mechanical and rheological properties (i.e., Young's modulus E, storage modulus G', loss modulus G'', and loss tangent tan(d)) of obtained hydrogels. Alginate-hydrogels stiffness is proportional to crosslinking density, polymer concentration and molecular weight of polymers. Hydrogels were physically crosslinked using different concentration of $CaCl_2$ (i.e., 0.1 M, 0.3M) to modulate stiffness.

| Hydrogel ID | Young's Modulus, E (Pa± SD) | Storage Modulus, G' (Pa mean ± SD) | Loss Modulus, G" (Pa mean ± SD) | Loss Tangent, tan(delta) (mean ± SD) |
|---|---|---|---|---|
| A1 | 5593 ± 562 | 4143 ± 484 | 332 ± 31 | 0.080 ± 0.006 |
| A3 | 12614 ± 1346 | 10293 ± 1457 | 1162 ± 190 | 0.113 ± 0.008 |
| B1 | 3042 ± 521 | 1283 ± 422 | 141 ± 31 | 0.118 ± 0.026 |
| B3 | 6598 ± 725 | 4151 ± 811 | 489 ± 105 | 0.111 ± 0.009 |

### SI.5 Optimization of cell culture media for co-culture

PC-3 cells and hTERT PF179T CAFs were co-cultured using optimized conditions. For co-culture cells were maintained in F-12:EMEM medium (1:1 volume ratio). EMEM medium was used without L-glutamine and supplements were reduced 50% for all supplements detailed earlier for respective single cell culture. This selection of medium is optimized to maintain cell proliferation and viability for both cell lines **(Figure SI.3)**.

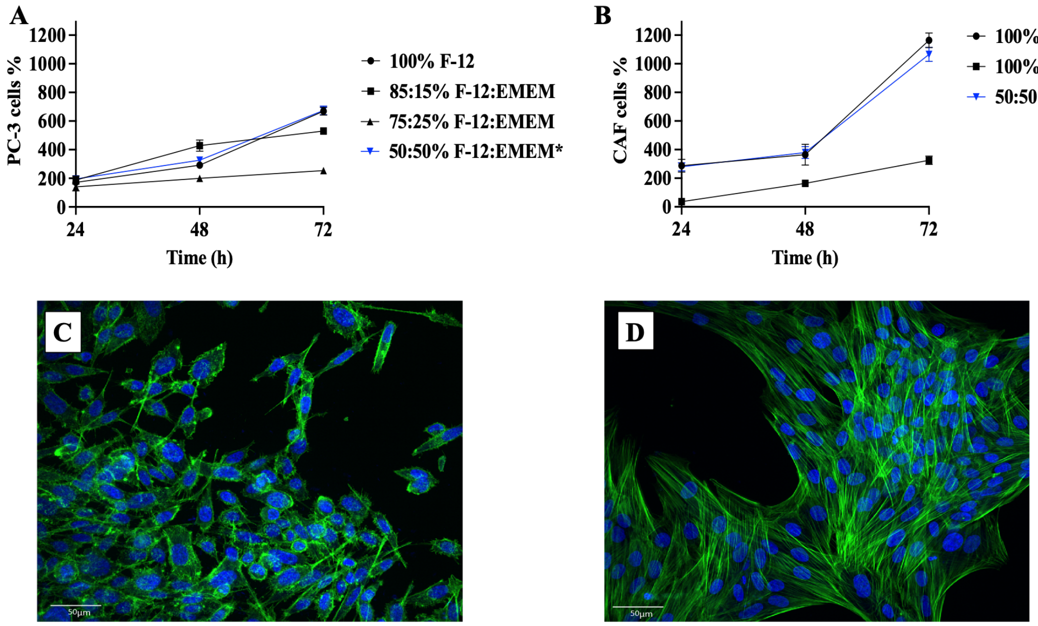


**Figure SI.3. Optimization of co-culture growth medium of co-culturing conditions**. Growth curve in different cell culture media at different ratio of F-12 and EMEM: **A)** PC-3 cells growth curve showing reduced growth curves when F12 is mixed with EMEM and compared to PC-3 cells proliferation in complete F-12 (used as control); **B)** hTERT PF179T CAF cells growth curve at different medium ratios, comparable CAFs proliferation is observed among complete EMEM (control) and 50:50% F-12:EMEM, with a reduced proliferation observed in complete F12 cell culture medium. Immunofluorescent images of: **C)** PC-3 and **D)** hTERT PF179T CAF cells. Cells were stained with DAPI (nuclei, blue) and Phalloidin (F- Actin, green), to evaluate cell morphology when cultured in the selected 50:50% F-12:EMEM. Of note, morphological variations where not observed up to 72 hours. Scale bars (50 μm).

### SI.6 Microbeads preparation and characterization for prostate- and stroma-specific microbeads

Preparation of prostate- and stroma-specific microbeads is sketched in **Figure SI.4**, while brightfield images are shown in (**Figure SI.5**). Semi-quantitative assessment of PC-3 cell viability was performed using Live/Dead assay (**Figure SI.6**). A small percentage of dead cells was measured at day 1 (~20-30%), with approx. 75-85% of live cells measured after 1 week of culture, in all models and confirming the metabolic activity measured.

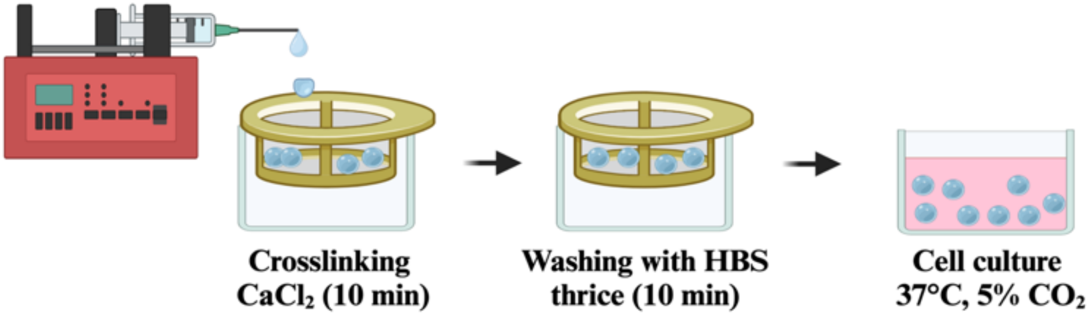


**Figure SI.4. Schematic representation of 3D *in vitro* models of PCa cells and CAFs in prostate-specific alginate hydrogels.** Hydrogels were prepared (as reported in the main manuscript, Table 1). Prior gelation with $CaCl_2$ (aq.), a homogenous suspension of cells (PC-3, hTERT PF179T CAF) was gently re-suspended in alginate-based solutions at a concentration of $1\times10^6$ cells/mL. By using a 27G.0.75" needle the cell suspension is ejected into a crosslinking solution of $CaCl_2$ (aq.) incubated for 10 min at RT allowing crosslinking. Obtained beads were washed in HBS thrice for 5 min each time and transferred in a well plate with complete cell culture media and incubated at 37°C, 5% $CO_2$. Created with Biorender.

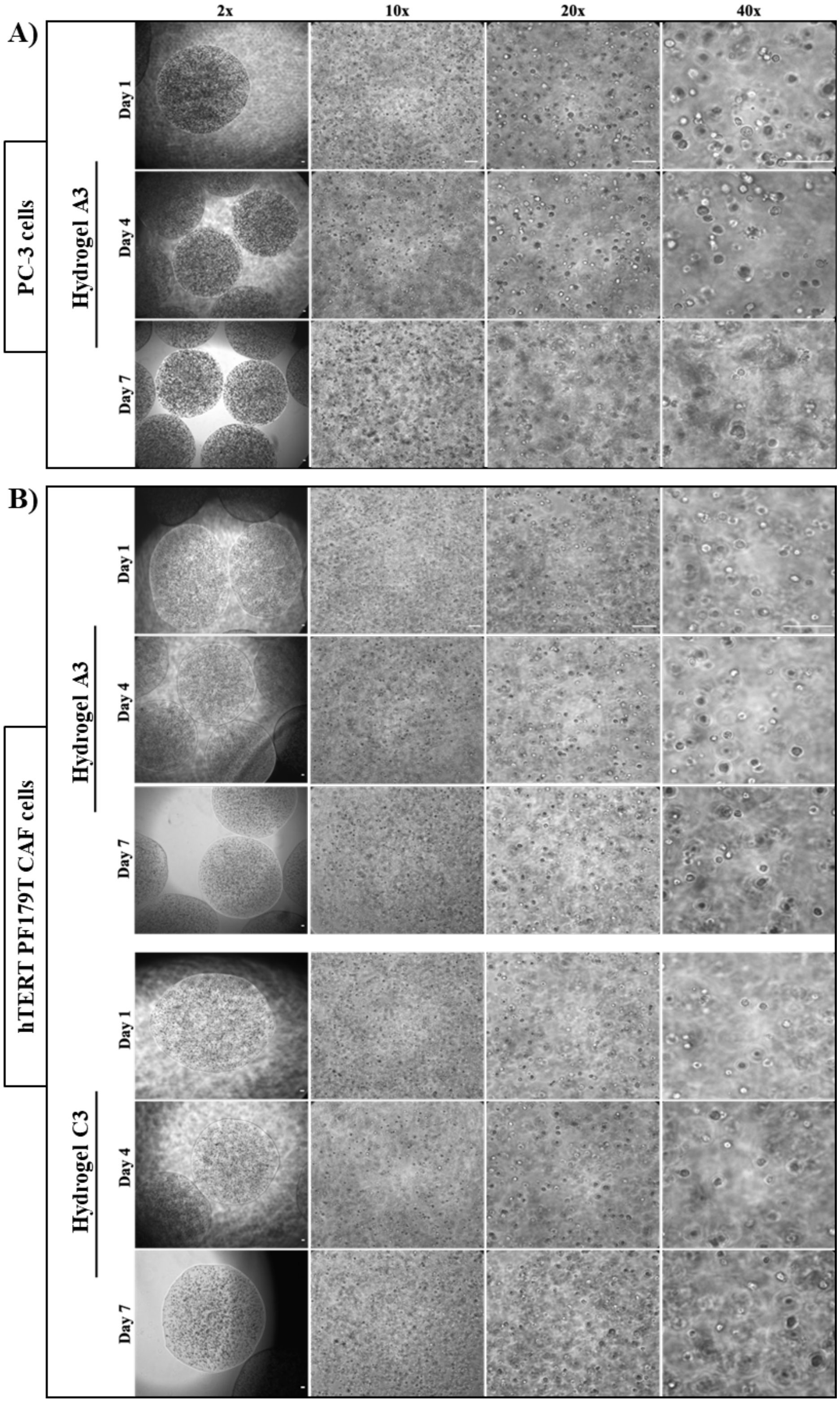


**Figure SI.5. Brightfield images of microbeads showing A)** PC-3 prostate 3D *in vitro* models and **B)** Stroma-specific 3D *in vitro* models**.** Representative images of PC-3 cells encapsulated in hydrogel **A3**, showing formation of larger PC-3 cell aggregates in the latter. Images acquired with a 2×, 10×, 20×, and 40× objective respectively, at day1, 4, and 7. Scale bars 100 μm.

Cells encapsulated in hydrogels **C1** and **C3** showed more uniformed and compact cellular aggregates compared to hydrogels **A1** and **A3**, indicating a higher level of reorganization and mobility within prostate-specific hydrogels at comparable stiffness in presence of PEG **(Figure SI6)**.

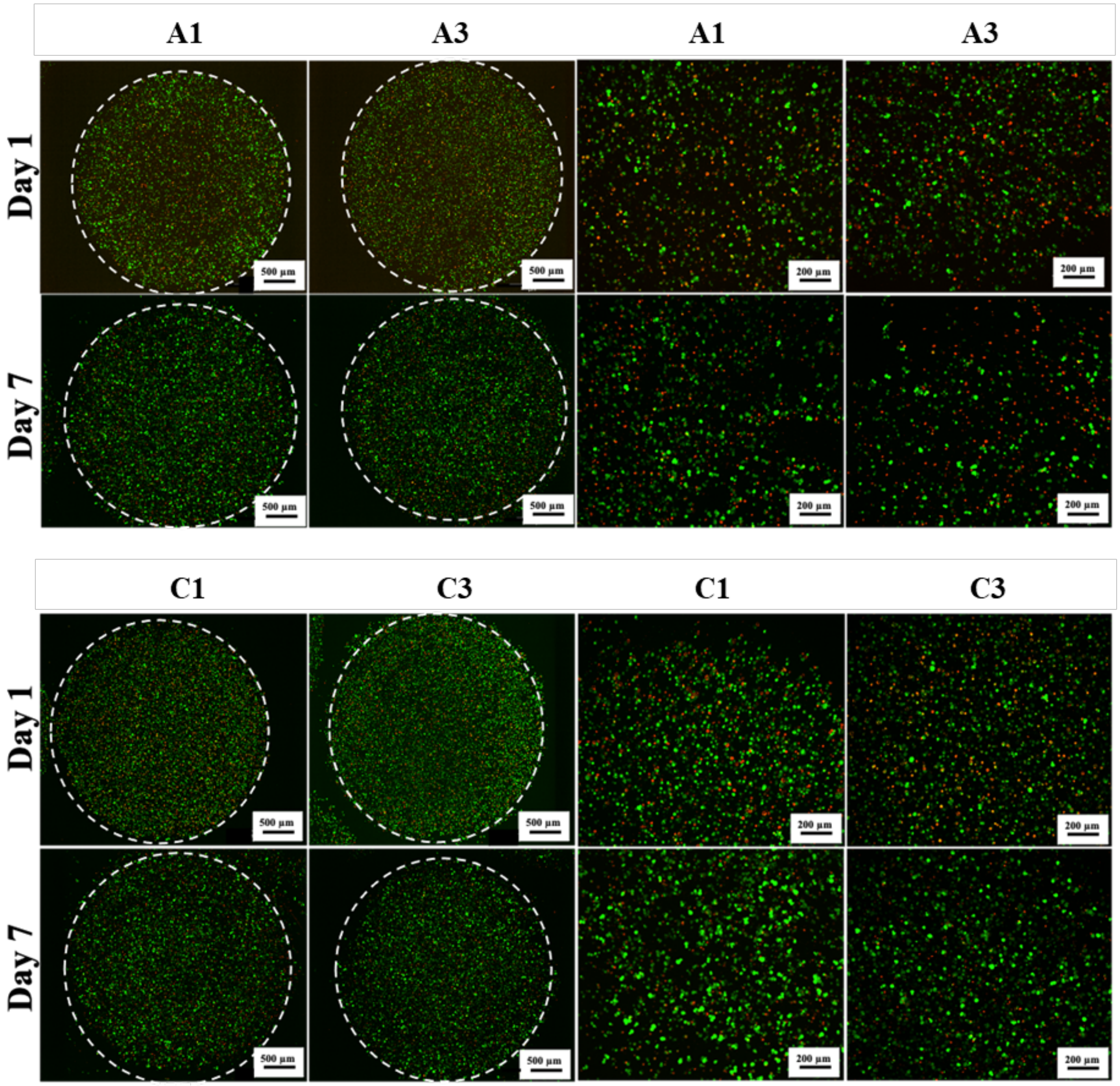


**Figure SI.6. Live/dead assay of PC-3 cells encapsulated in different prostate-specific hydrogels.** Fluorescent images at different magnification show live proliferative cells (green cytoplasm) compared to dead cells (red nuclei) observed at different time points (i.e., day 3, day 7). Scale bars: 500 µm and 200 µm.

### SI.7 Migration assay: PC-3 migratory phenotype after conditioning in hydrogel C1 and C3

PC-3 cells preconditioned (7 days) in *prostate-specific microbeads* were recovered and seeded in 24-well plate at a density of $1.5\times10^5$ cells/well and routinely cultured allowing cell adhesion (48 h, 37°C, 5% $CO_2$). A scratch was then performed in each well using a sterile 200 µL tip, cellular debris was removed by gentle washing with cell culture media to avoid cellular detachment. Cells were then cultured in low serum media (1% v/v FBS in F-12 with 1% v/v L-glutamine and 1% v/v PenStrep) to maintain low cell proliferation. Brightfield images for scratch assay were acquired after 0, 24, and up to 120 h using an inverted microscope (Leica DMI6000, Leica Microsystems, UK). The area of scratch invaded by cells was calculated using ImageJ (v1.53a) by measuring the combined cellular area in the scratch over time.

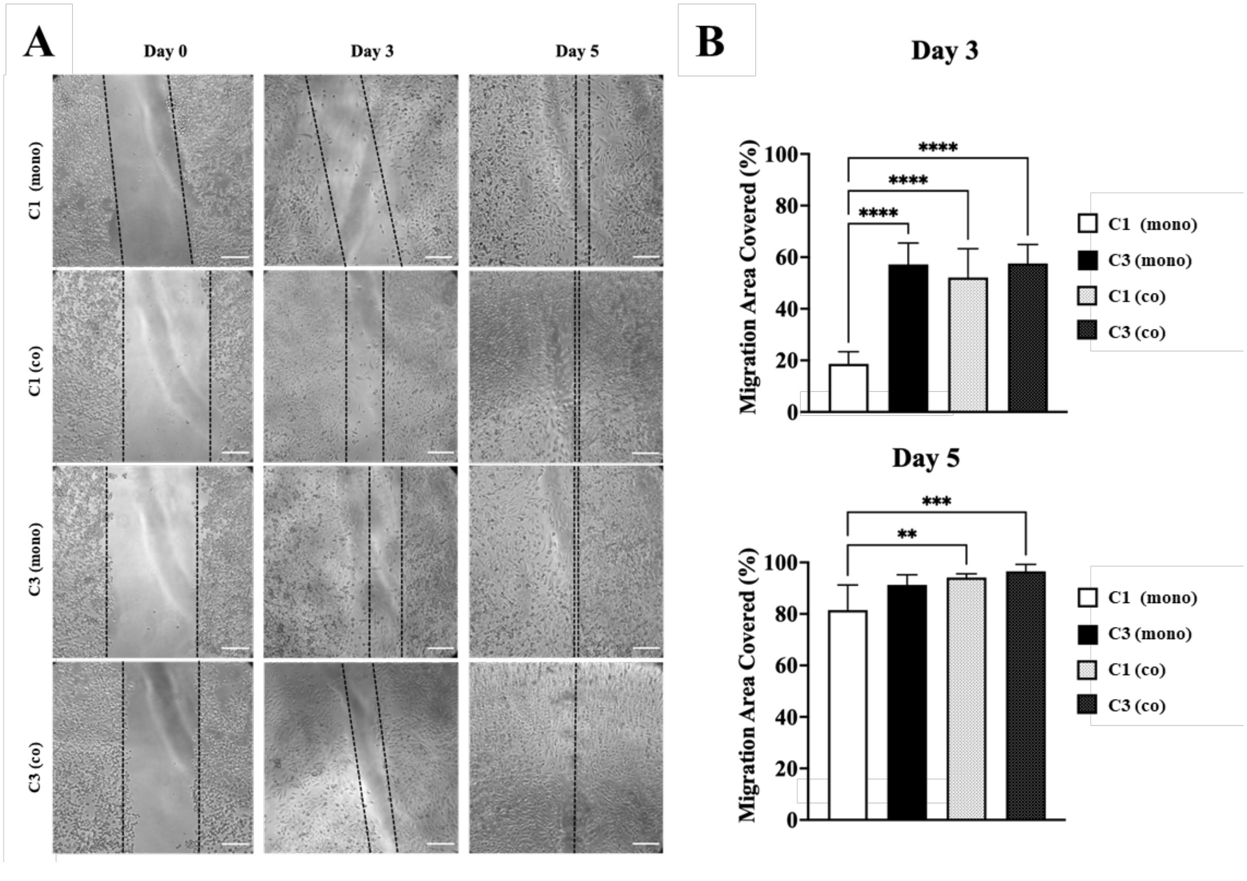


**Figure SI.7. Scratch assay of PC-3 cells PC-3 conditioned in engineered 3D prostate in vitro models. A)** Brightfield images of the scratch assay, cells were monitored for 5 days. (Scale bars 200 µm). **B)** Graphical representation of scratch area (%) covered by migratory PC-3 cells in µm$^2$, data is represented as mean and SD of N=2, n=2 independent experiments; P-values represented as *$p \leq 0.05$, **$p \leq 0.01$, ***$p \leq 0.001$, ****$p \leq 0.0001$).

**SI.8 Invasion assay: PC-3 migratory phenotype after conditioning in prostate-specific hydrogels**

PC-3 cells preconditioned (7 days) in *prostate-specific microbeads* were recovered, stained with Cytopainter red by incubation with 1× dye diluted in cell culture media (1 h, 37°C, 5% $CO_2$). Cells were then washed trice with HBS and then embedded in a 1 mg/mL collagen hydrogel precursor diluted in 10× PBS and sterile water, adjusted to pH ~7.4 using sterile 1 M NaOH solution (aq.) and 7.5% NaHCO3 solution (aq.) following the supplier's instruction. Cell-laden hydrogels were routinely cultured (37°C, 5% $CO_2$) and the invasion imaged at day 0 and day 3 with confocal microscope using (Ex/Em 570/600 nm) filter to detect Cytopainter (live cells, red).

To ensure monitoring of PC-3 cells preconditioned in engineered 3D prostate *in vitro* models, cells were stained (i.e., Cytopainter red) and calls invasion into collagen hydrogels monitored over time, following our previous study [2].

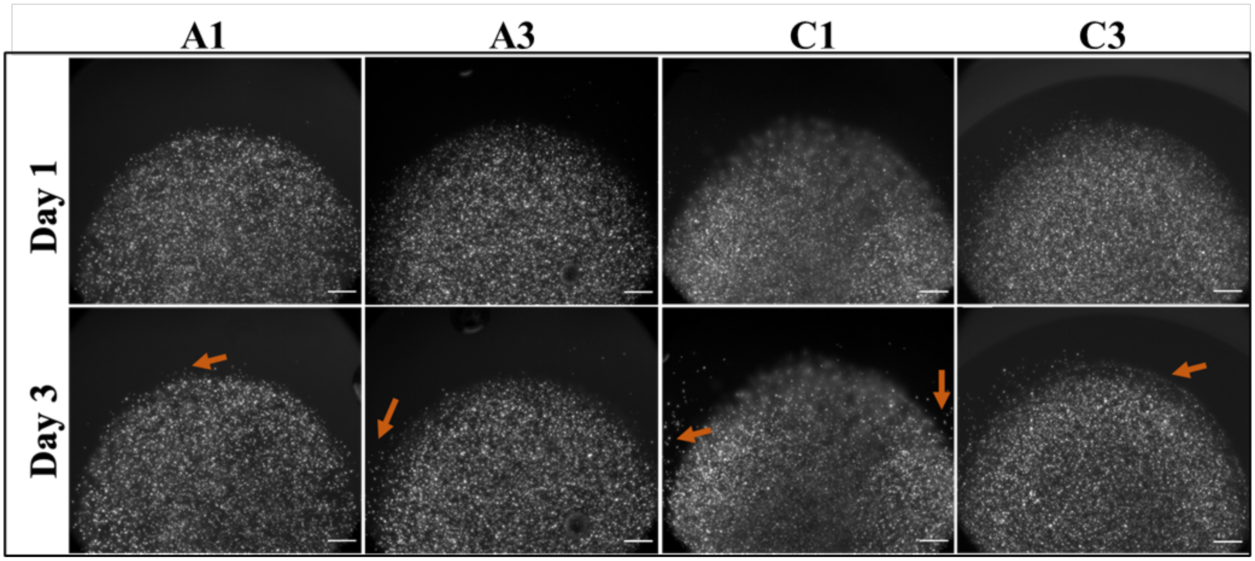


**Figure SI.8. Invasion ability of PC-3 conditioned in engineered 3D prostate in vitro models.** Immunofluorescence images of Cytopainter red stained PC-3 cells then embedded in collagen hydrogel. Cells were imaged at day 0 and day 3 to evaluate invasion after pre-conditioning in four different hydrogels varying in composition and viscoelastic properties. (Scale bars 100 µm).

### SI.9 Assessment of invasive phenotype of PC-3 cells

Expression of PCa progression and EMT was observed in PC-3 cells after 3 days of culture on standard *in vitro* models. Data were used as control to evaluate the impact of the 3D microenvironment on PC-3 phenotypes. The same protocol for staining and detection reported in the main manuscript was used.

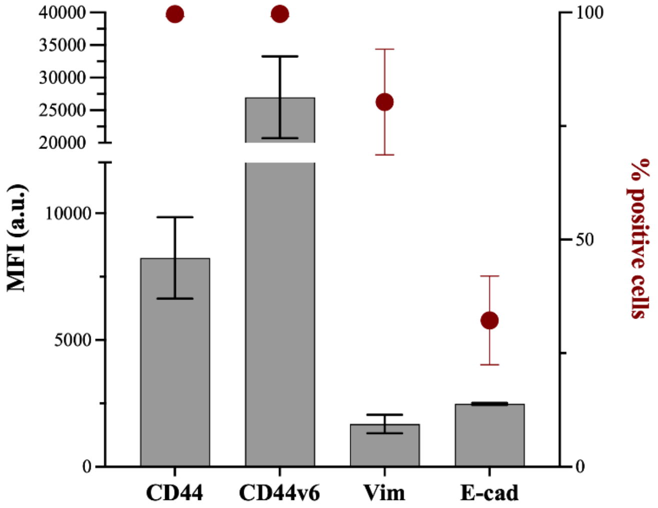


**Figure SI.9. Flow cytometry analysis of EMT markers and CD44.** Flow cytometry quantification in PC-3 cells after 3 days culture in 2D.
